# Large library docking for cannabinoid-1 receptor agonists with reduced side effects

**DOI:** 10.1101/2023.02.27.530254

**Authors:** Tia A. Tummino, Christos Iliopoulos-Tsoutsouvas, Joao M. Braz, Evan S. O’Brien, Reed M. Stein, Veronica Craik, Ngan K. Tran, Suthakar Ganapathy, Fangyu Liu, Yuki Shiimura, Fei Tong, Thanh C. Ho, Dmytro S. Radchenko, Yurii S. Moroz, Sian Rodriguez Rosado, Karnika Bhardwaj, Jorge Benitez, Yongfeng Liu, Herthana Kandasamy, Claire Normand, Meriem Semache, Laurent Sabbagh, Isabella Glenn, John J. Irwin, Kaavya Krishna Kumar, Alexandros Makriyannis, Allan I. Basbaum, Brian K. Shoichet

**Affiliations:** Department of Pharmaceutical Chemistry, University of California, San Francisco, San Francisco, CA 94158, USA; Graduate Program in Pharmaceutical Sciences and Pharmacogenomics, University of California, San Francisco, San Francisco, CA 94158, USA; Center for Drug Discovery and Department of Pharmaceutical Sciences, Northeastern University, Boston, MA 02115, USA; Department of Anatomy, University of California, San Francisco, San Francisco, CA 94158, USA; Department of Molecular and Cellular Physiology, Stanford University School of Medicine, Stanford, CA 94305, USA; Division of Molecular Genetics, Institute of Life Science, Kurume University, Fukuoka, Japan; Enamine Ltd., 67 Chervonotkatska Street, Kyiv, 02094, Ukraine; National Taras Shevchenko University of Kyiv, 60 Volodymyrska Street, Kyiv 01601, Ukraine; Chemspace LLC, 85 Chervonotkatska Street, Suite 1, Kyiv, 02094, Ukraine; National Institute of Mental Health Psychoactive Drug Screening Program (NIMH PDSP), School of Medicine, University of North Carolina at Chapel Hill School of Medicine, Chapel Hill, NC 27599, USA; Domain Therapeutics North America Inc., Montréal, Québec, H4S 1Z9, Canada, Montréal, QC, H3T 1J4, Canada; Department of Chemical and Chemical Biology, Northeastern University, Boston, MA 02115, USA

## Abstract

Large library docking can reveal unexpected chemotypes that complement the structures of biological targets. Seeking new agonists for the cannabinoid-1 receptor (CB1R), we docked 74 million tangible molecules, prioritizing 46 high ranking ones for *de novo* synthesis and testing. Nine were active by radioligand competition, a 20% hit-rate. Structure-based optimization of one of the most potent of these (K_i_ = 0.7 µM) led to **‘4042**, a 1.9 nM ligand and a full CB1R agonist. A cryo-EM structure of the purified enantiomer of **‘4042** (**‘1350**) in complex with CB1R-G_i1_ confirmed its docked pose. The new agonist was strongly analgesic, with generally a 5-10-fold therapeutic window over sedation and catalepsy and no observable conditioned place preference. These findings suggest that new cannabinoid chemotypes may disentangle characteristic cannabinoid side-effects from their analgesia, supporting the further development of cannabinoids as pain therapeutics.

## INTRODUCTION

Although the therapeutic use of cannabinoids dates back to at least the 15^th^ century^1,2^, their use in modern therapy, for instance as analgesics, has been slowed by their sedative and mood-altering effects, and by concerns over their reinforcing and addictive potential^3,4^. With changes in cannabis’ legal status, an ongoing epidemic of chronic pain, and efforts to reduce reliance on opioids for pain management, has come a renewed interest in understanding both the endocannabinoid system and how to leverage it for therapeutic development^5^. Areas of potential application of such therapeutics include anxiety^6^, nausea^7^, obesity^8^, seizures^9^, and pain^10^, the latter of which is the focus of this study. Progress has been slowed by the physical properties of the cannabinoids themselves, which are often highly hydrophobic, by the challenges of the uncertain legal environment, and by the substantial adverse side effects often attending on the drugs, including sedation, psychotropic effects, and concerns about reinforcement and addiction^3^. Indeed, a characteristic defining feature of cannabinoids is their “tetrad” of effects^11^: analgesia, hypothermia, catalepsy, and hypolocomotion, the latter three of which may be considered adverse drug reactions. Meanwhile, inconclusive results in human clinical trials^12^ have led to uncertainty in the field as to the effectiveness of cannabinoids as therapeutics. Nevertheless, the strong interest in new analgesics, and the clear efficacy of cannabinoids in animal models of nociception^13^, have maintained therapeutic interest in these targets.

The cannabinoid-1 and -2 receptors (CB1R and CB2R), members of the lipid family of G-protein coupled receptors (GPCRs), are the primary mediators of cannabinoid activity^14^. The structural determination of these receptors^15–21^ affords the opportunity to use structure-based methods to find ligands with new chemotypes. Recent structure-based docking of make-on-demand virtual libraries have discovered new chemotypes for a range of targets, often with new pharmacology and reduced side effects^22–29^. Thus, new CB1R chemotypes emerging from a structure-based approach might address some of the liabilities of current cannabinoids, such as their physicochemical properties or side-effect profiles. Seeking such new chemotypes, we computationally docked a library of 74 million virtual but readily accessible (“tangible”) molecules against CB1R, revealing a range of new scaffolds with relatively favorable physical properties. Structure-based optimization led to agonists binding with low-nanomolar affinities. The lead agonist is a potent analgesic, with pain-relieving activity at doses as low as 0.05 mg/kg. It has a five to ten-fold separation between analgesia and both sedation and catalepsy, addressing two of the four aspects of the “tetrad” and highlighting the utility of large library structure-based screens for identifying new pharmacology through new chemical scaffolds.

## RESULTS

### Large-library docking against CB1R

The CB1R orthosteric site is large and lipophilic, explaining the high molecular weight and hydrophobicity of many of its ligands (**Fig. S1**); these physical properties are often metabolic and solubility liabilities^30^. We therefore sought molecules in a more “lead-like” physical property range. In preliminary studies, strict enforcement of such properties (MW ≤ 350 amu, cLogP ≤ 3.5) led to no new ligands from docking. We therefore focused on a 74-million molecule subset of the ZINC15 database^31^ composed of molecules between 350 and 500 amu with calculated LogP (cLogP) of between 3 and 5, reasoning that these would be more likely to complement the CB1R site, while retaining polarity and size advantages over many cannabinoids (**Fig. 1B**). Each molecule was docked in an average of 3.04 million poses (orientations x conformations), totaling 63 trillion sampled and scored complexes. Seeking a diverse set of candidates, the top-ranking 300,000 were clustered into 60,420 sets and the highest scoring member of each was filtered for topological dissimilarity to known CB1/CB2 receptor ligands in ChEMBL^32,33^ using Tanimoto coefficient (Tc < 0.38) comparisons of ECFP4-based molecular fingerprints. High-ranking compounds that did not resemble known ligands were filtered for potential polar interactions with S383^7.39^ and H178^2.65^ (superscripts denote Ballesteros-Weinstein nomenclature^34^; see **Methods, Fig. 1A, Supplementary Table 1**). The top-ranking 10,000 remaining molecules were visually evaluated in UCSF Chimera^35^, and 60 were prioritized for *de novo* synthesis. Of these, 46 were successfully made and tested for CB1R activity. Consistent with the design of the library, the new molecules were smaller and more polar than most existing cannabinoid ligands, skirting the edge of property-space that is suitable for the large and hydrophobic CB1R orthosteric pocket (**Fig. 1B**).

**Figure 1.**
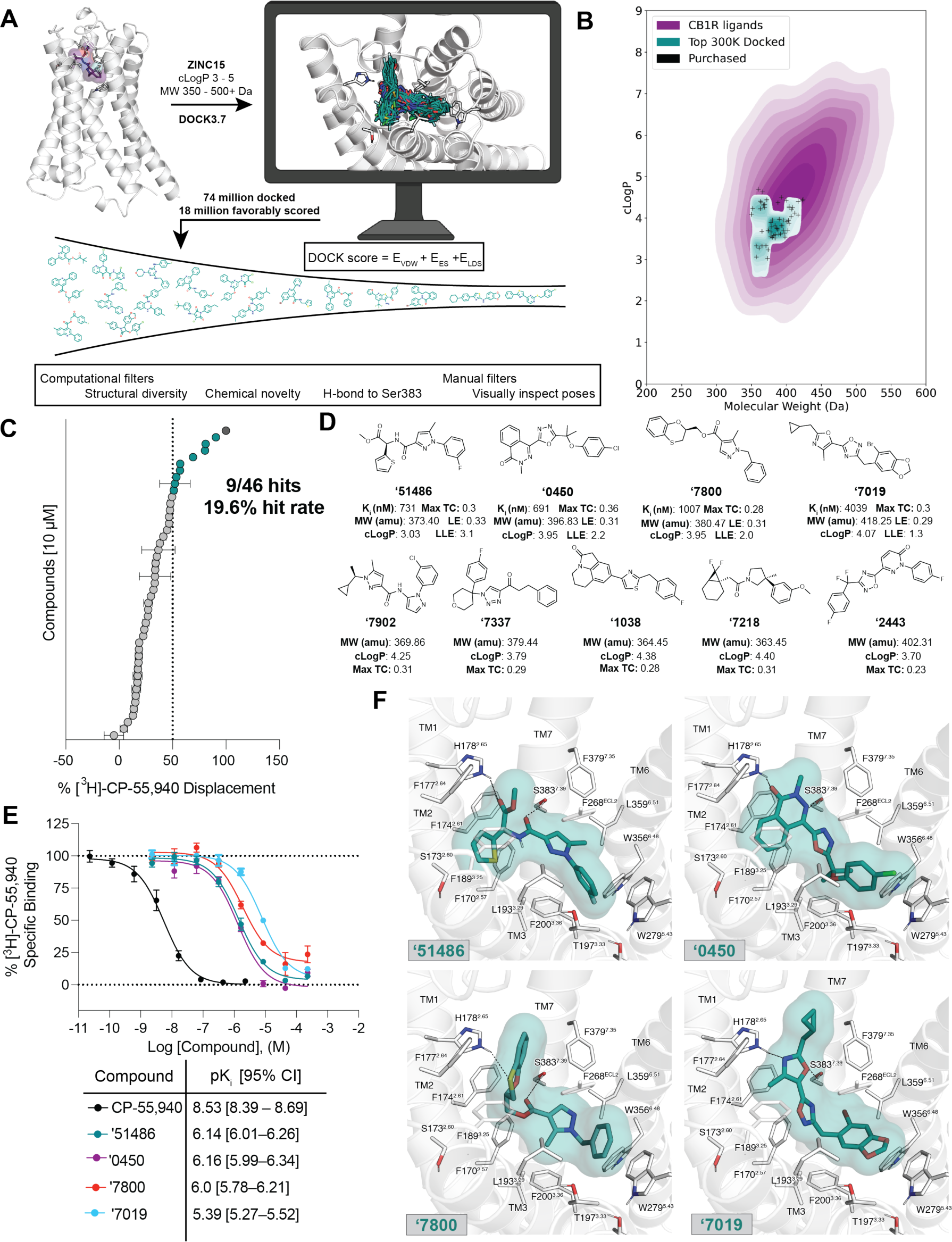
Large-scale docking of a 74-million molecule library against the CB1R. **A.** Workflow of the docking campaign. **B.** Overlap of physical properties of CB1R ligands versus the top docked and purchased ligands. **C.** Single-point radioligand displacement data for the 46 tested compounds. **D.** 2D structures and properties of the nine hits. **E.** Secondary binding assay for the top four hits. **F.** Docked poses of the top four hits with H-bonds and other binding pocket residues indicated. Data in panels **C.** and **E.** represent mean ± SEM from three independent experiments.

In single-point radioligand displacement experiments, 9 of the 46 prioritized molecules displaced over 50% of the radioligand, a 20% hit-rate (**Fig. 1C-D**, **Supplementary Table 1**). The top four of these (ZINC537551486, ZINC1341460450, ZINC749087800, and ZINC518437019, referred to as **‘51486**, **‘0450**, **‘7800**, and **‘7019**, respectively, from here on) were then tested in full concentration-response. All four displaced the radioligand ^3^H-CP-55,940, with K_i_ values ranging from 0.7 to 4 µM (**Fig. 1E**). Owing to coupling to the inhibitory G_αi_ G-protein, functional efficacy experiments monitoring a decrease in forskolin (FSK) simulated cAMP were tested using hCB1-expressing cells, with **‘51486** and **‘0450** showing modest agonism. Limited solubility prohibited testing at high enough concentrations to obtain accurate EC_50_ values; fortunately, colloidal aggregation counter-screens showed no such activity below 10 µM (**Fig. S2**), suggesting that activity seen in binding and functional assays is not due to this confound^36^. Taken together, the nine actives explore a range of chemotypes topologically unrelated to known CB1 ligands (**Supplementary Table 1**), with relatively favorable physical properties (**Fig 1B,D**).

Although the new ligands are chemically and physically distinct from established cannabinoids, their docked poses recapitulate the interactions of the known ligands but do so with different scaffold and recognition elements. All of the four most potent ligands docked to adopt a “C” shaped conformation characteristic of the experimentally observed geometries of MDMB-Fubinaca^18^, AM11542, and AM841^16^ bound to CB1R. Similarly, all four are predicted to hydrogen-bond with S383^7.39^, a potency-determinant interaction at CB1 receptors observed in nearly all agonist-bound ligand-receptor complexes^37,38^. Additionally, all four ligands are predicted to make secondary hydrogen bonds to H178^2.65^, a feature thought to be important for potency as well as agonism of CB1R^38^. Largely, these electrostatic interactions are made using unique hydrogen-bond acceptor groups, such as an oxazole, oxathiine, or pyridazinone. Other characteristic hydrophobic and aromatic stacking interactions are found throughout the ligands, including with F268^ECL2^, W279^5.43^, and F174^2.61^, though again often using different aromatic groups than found in the known ligands (**Fig. 1F**). Similarly, all four ligands exhibit aromatic stacking and hydrophobic packing with the twin-toggle switch residues W356^6.48^ and F200^3.36^ which are important for receptor activation^39,40^.

We sought to optimize these initial ligands. Molecules with ECFP4 Tcs ≥ 0.5 to the four actives were sought among a library of 12 billion tangible molecules using SmallWorld (NextMove Software, Cambridge UK), a program well-suited to ultra-large libraries. These analogs were built, docked, filtered, and selected using the same criteria as in the original docking campaign. Between 11 and 30 analogs were synthesized for each of the four scaffolds. Optimized analogs were found for three of the four initial hits, improving affinity by between 5 and 24-fold, with **‘51486** improving 16-fold to a K_i_ of 44 nM, **‘7019** improving 5-fold to 87 nM, and **‘0450** improving 24-fold to 163 nM (**Supplementary Table 2**). In subsequent bespoke synthesis, the 44 nM analog of **‘51486**, **‘60154**, was further optimized to compound Z8504214042 (from here on referred to as **‘4042**) with a K_i_ of 1.9 nM (**Fig. S3**). **Figure 2** summarizes the structure-activity relationship (SAR) of the **‘51486/‘4042** series.

**Figure 2.**
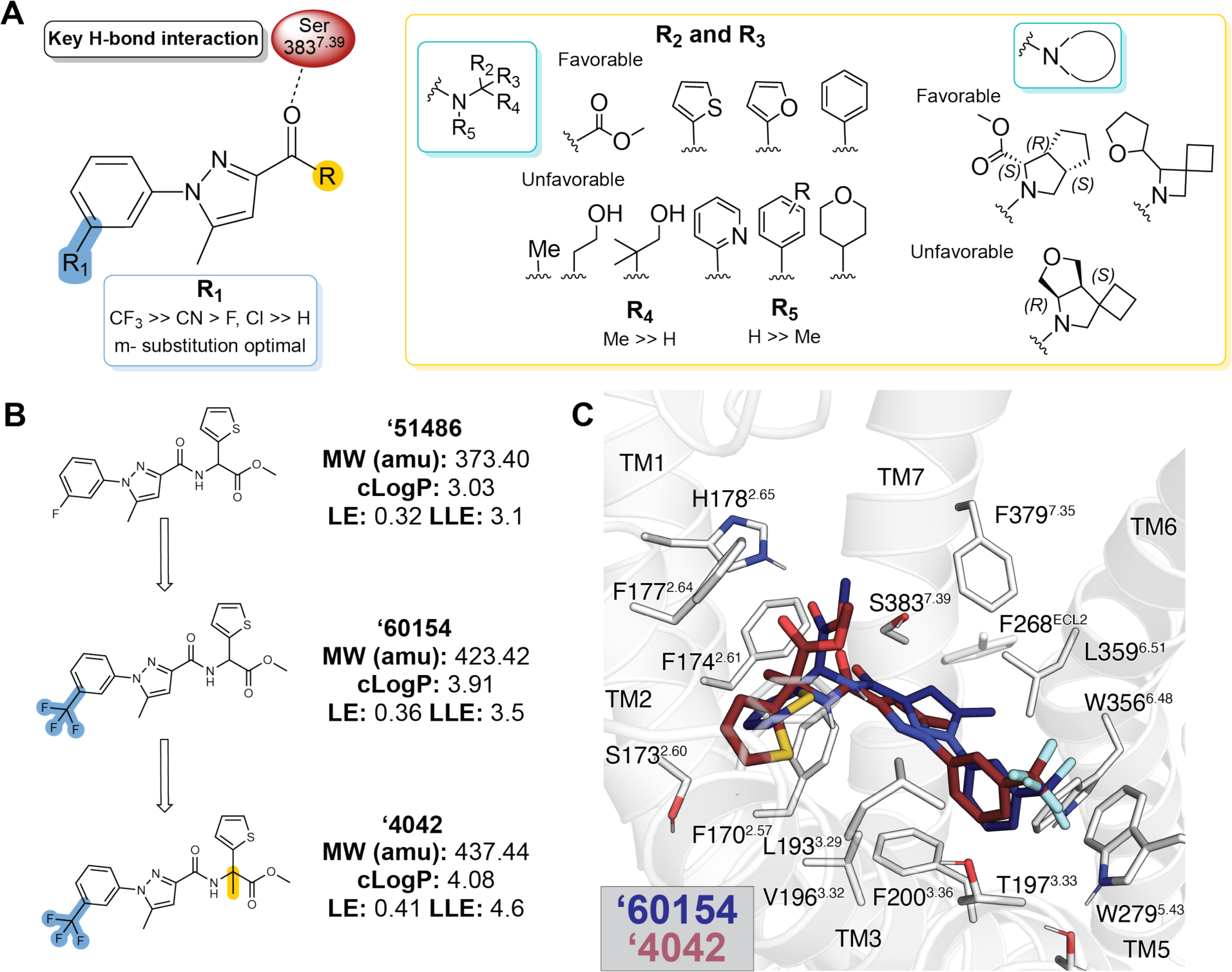
Structure-activity relationships and optimization of ‘51486 to ‘4042. **A.** Pharmacophore model based on the structure-activity relationships discovered via analoging ‘**51486**. **B.** 2D structures of the docking hit **‘51486** and analogs that lead to **‘4042**. **C.** Docking predicted pose of **‘60154** (navy) and **‘4042** (purple).

Key learnings from the SAR include the importance of bulky and hydrophobic groups in the R_1_ position of **‘4042**, which is modeled to pack against W279^5.43^ and T197^3.33^ and methylation of the chiral center (R_4_ position), which is predicted to increase van der Waals interactions between the ligand and transmembrane helix 2. Finally, the terminal ester is modeled to hydrogen-bond with H178^2.65^ of the receptor, though the distance suggests either a water-mediated interaction, or a weak hydrogen bond. As expected, the carboxylate analog of the ester, **‘4051**, bound only weakly (K_i_ = 5 µM, 5,000-fold less potent)—this molecule, a very close analog to **‘4042**, may provide the inactive member of a “probe pair” for future research. The lead that emerged, **‘4042** at 1.9 nM, is about 2-fold more potent than the widely used CB1R probe CP-55,940 (**Fig. 4B**, below) and equipotent to the marketed drug nabilone (**Fig. S3A**, **Supplementary Table 2**). Although more hydrophobic than the initial docking hit ‘**51486**, its lipophilic ligand efficiency improved from 3.1 to 4.6 (**Fig. 2B**).

#### Cryo-EM structure of the ‘1350-CB1R-G_i1_ complex

To understand the SAR of the **‘4042** series at atomic resolution, and to template future optimization, we determined the structure of the agonist in complex with the activated state of the receptor. Initial efforts at single particle cryo-electron microscopy (cryo-EM) of **‘4042** in complex with CB1R and the G_i1_ heterotrimeric G-protein led to a structure where the ligand density seemed to reflect either multiple conformations of a single ligand, or multiple ligands. As **‘4042** is a racemate, we purified it into it its component isomers, **‘1350** and **‘8690** using chiral chromatography (**Fig. S4**) and measured CB1R binding by radioligand competition. With K_i_ values of 0.95 nM and 90 nM, respectively, **‘1350** was substantially more potent than its enantiomer, and subsequent functional studies revealed it to be the much stronger agonist (**Fig. 4A-B, Fig. S4**; below). Accordingly, we determined the cryo-EM structure of the **‘1350-CB1R-G_i1_** complex (**Fig. 3**, **Fig. S5,** see **Methods**) to a nominal resolution of 3.3 Å (**Supplementary Table 3**). Consistent with earlier structures of CB1R in its activated state, the ligand occupies the orthosteric pocket formed by transmembrane helices (TMs) 2-3 and 5-7 and is capped by extracellular loop (ECL) 2.

**Figure. 3.**
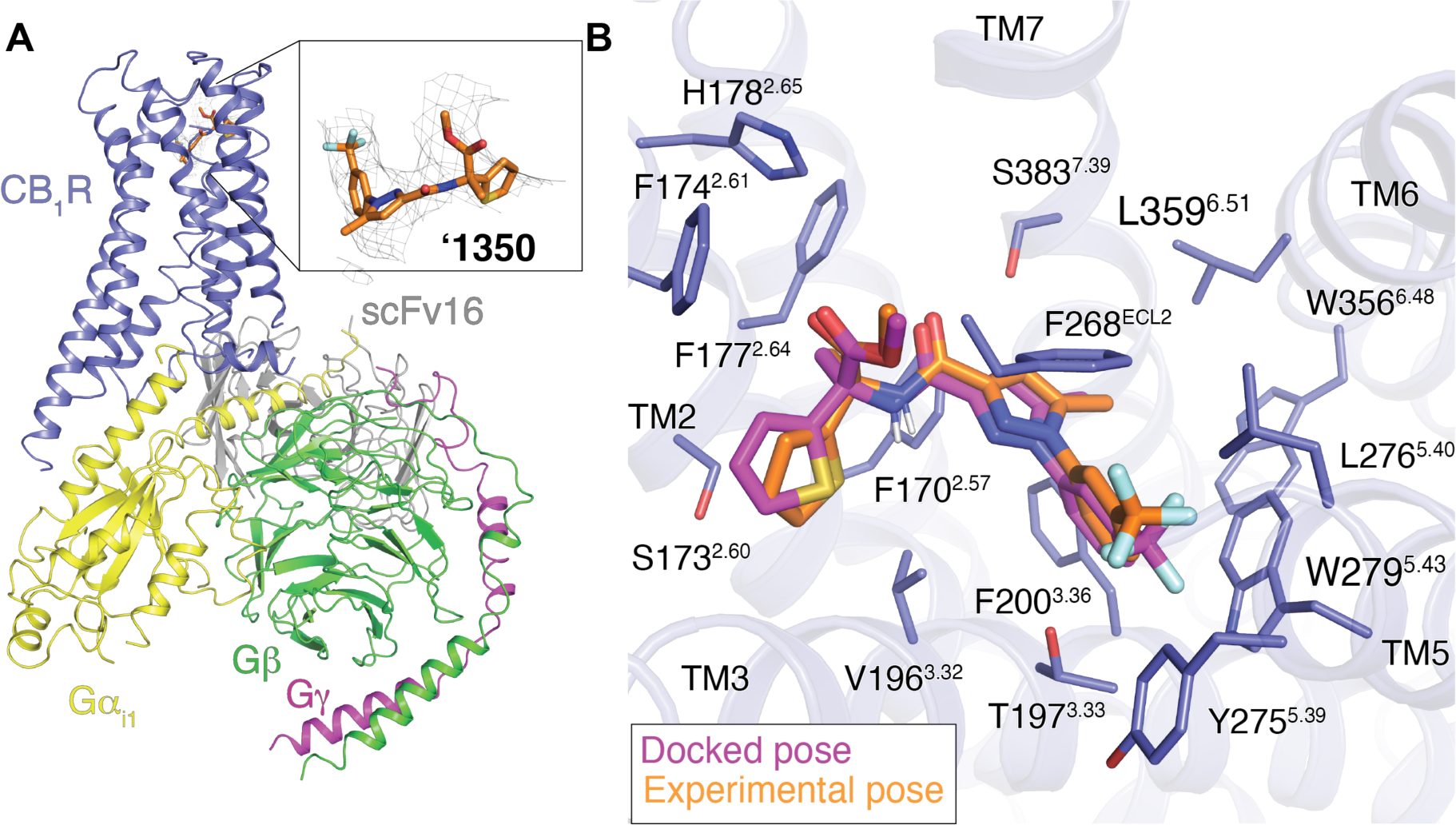
Cryo-EM structure of ‘1350-CB1R-Gi1 complex. **A.** Cryo-EM structure of **‘1350-CB1R-G_i1_** highlighting the ligand density. **B.** Overlay of the docked pose (magenta) with the experimental pose (orange) of **‘1350.**

The experimental structure of ‘**1350** superposes well on the docking-predicted pose of **‘4042** in its *R*-enantiomer, which was the enantiomer with the better docking score to the receptor (−43 DOCK3.7 score versus −38 DOCK3.7 score for the *S*-enantiomer). The predicted and experimental structures superposed with an all-atom RMSD of 1.1 Å (**Fig. 3B**). The major interactions with CB1R predicted by the docking are preserved in the experimental structure, including the key hydrogen-bond between the amide carbonyl of the ligand and S383^7.39^. The trifluoromethyl group is complemented by van der Waals and quadrupole interactions with residues W279^5.43^ and T197^3.33^, as anticipated by the docked structure, and consistent with the improvement in affinity by −1.7 kcal/mol (17-fold in K_i_) on its replacement of the original fluorine.

#### Agonism and subtype selectivity of ‘4042

Given the potent affinity of **‘4042** and of **‘1350** (**Fig. 4A**), we next investigated their functional activity, and how they compared to that of the widely studied cannabinoid, CP-55,940^2^. We first measured G_i/o_ mediated agonism via inhibition of forskolin-stimulated cAMP in the Lance Ultra cAMP assay (**Methods**). Both **‘4042**, **‘1350**, and several of its analogs are agonists in human CB1R-expressing cells (hCB1R), with EC_50_ values commensurate with their affinities (**Supplementary Table 2,4 Fig. S3, S6-7**) and with efficacies close to full agonism (E_max_ typically > 75%). **‘4042** and **‘1350** had hCB1R EC_50_ (E_max_) values of 3.3 nM (78%) and 1.6 nM (77%) (**Fig. 4B**). The activity of racemic **‘4042** was confirmed in several orthogonal cAMP and ß-arrestin assays (see **Methods**), including in the Cerep cAMP assay (**Fig. S3C**), the Glosensor assay (**Fig. S3D**), the Tango ß-arrestin translocation assay (**Fig. S3E**) and the DiscoverX ß-arrestin-2 recruitment assay (**Fig. S3F**). In summary, **‘4042** and its *R*-isomer, **‘1350,** are potent agonists of hCB1R with low nM EC_50_ values.

**Figure 4.**
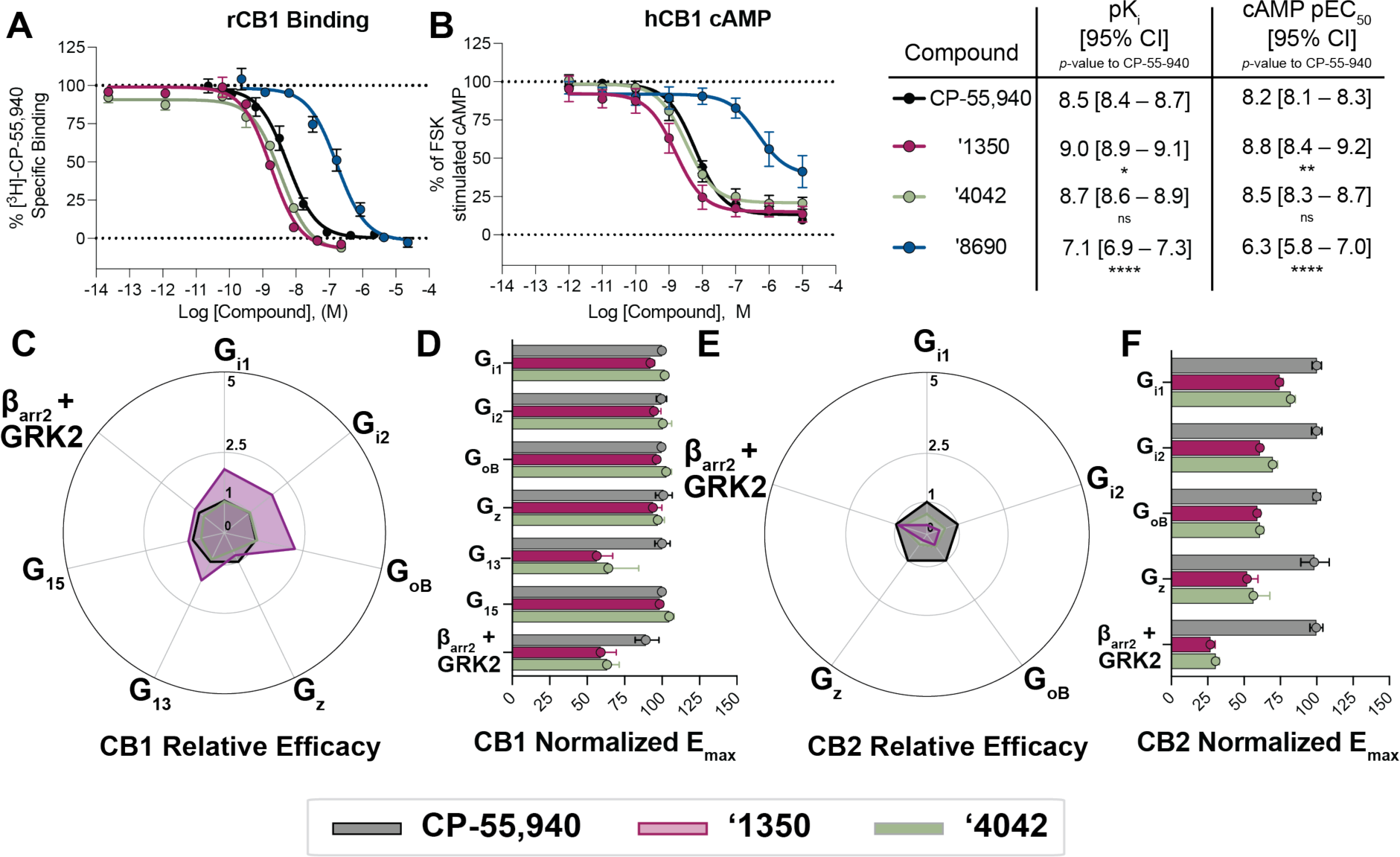
Functional activity of ‘4042 and its active enantiomer ‘1350. **A.** Binding affinity or **B.** Functional cAMP inhibition of **‘4042** and its enantiomers **‘1350** and **‘8690** compared to CP-55,940. One-way ANOVA statistical significance of individual pKi (**A**) or pEC50 (**B**) comparisons to CP-55,940 after correction with Dunnett’s test of multiple hypotheses are depicted in the table; ns = not significant, * p<0.05, ** p<0.01, **** p<0.001. **C.** Relative efficacy of **‘1350** and **‘4042** compared to CP-55,940 at hCB1. **D.** Normalized E_max_ from the experiments in **C. E.** Relative efficacy of ‘**1350** and **‘4042** compared to CP-55940 at hCB2. **F.** Normalized E_max_ from the experiments in **E.** Data in **A. & B.** represent mean ± SEM from three independent experiments. Data in **D & F.** represent mean ± 95% CI of the best-fit E_max_ value.

Fortified by this potent activity, and to control for system bias^41–43^ and questions of signal amplification in the cAMP assays, we investigated both **‘4042** and the more active of its stereoisomers, **‘1350**, for differential recruitment of several G-proteins and β-arrestin-2 against both CB1R and CB2R in the ebBRET bioSens-All^®^ platform, comparing its activity to CP-55,940 (**Fig. 4C-F, Fig. S6, Supplementary Table 5-6**). A useful way to picture the differential effects of **‘1350** and **‘4042** relative to CP-55,940 at CB1R and CB2R is via “radar” plots (**Fig. 4C** and **4E**) depicting the relative effectiveness^41^ toward each signaling pathway (10^Δlog(Emax/EC50)^, see **Methods**). In CB1R, **‘1350** was approximately 2 times more relatively efficacious at recruiting G_i/o_ and G_13_ subtypes than was CP-55,940, though the pattern of effectors recruited was similar. Similar coupling profiles were seen for **‘4042**, though the effects were smaller, consistent with the latter compound being an enantiomeric mixture. Whereas the CB1R radar plots were similar in pattern for **‘1350**, **‘4042** and CP-55,940, the differential activities for the highly related CB2R differed qualitatively (**Fig. 4E-F; Fig. S6; Supplementary Table 7-8**). Although the affinity of **‘4042** at the two receptors is almost undistinguishable (**Fig. S8**), there was a marked difference in functional activity, with **‘4042** consistently being a weaker efficacy partial agonist at CB2R (**Fig. S6C-D, S8**) versus its essentially full agonism at CB1. This was true for the racemate **‘4042** as well as its active enantiomer **‘1350** across four separate functional assays including the bioSens-All^®^ BRET assay, the Lance Ultra cAMP assay, TRUPATH BRET2 assay, and the Tango β-arrestin recruitment assay (**Fig. S8B-D**). Indeed, whereas against CB1R **‘1350** had greater relative efficacy against inhibitory G-proteins versus CP-55,940, in CB2R the pattern was reversed, with CP-55,940 being substantially more relatively efficacious than **‘1350** (**Fig. 4C-F**).

### The new CB1R agonist is analgesic with reduced cannabinoid side effects

#### Off-target selectivity and pharmacokinetics

Encouraged by the potency and functional selectivity, and the negligible functional differences between the racemic and enantiomeric mixture, we progressed **‘4042** into *in vivo* studies for pain relief. We began by investigating the selectivity of **‘4042** against potential off-targets. **‘4042** was tested first for binding and functional activity against a panel of 320 GPCRs and 46 common drug targets at the PDSP (**Fig. S9**). Little activity was seen except against the melatonin-1 (MT1R), ghrelin (GHSR), Sigma1 and peripheral benzodiazepine receptors. In secondary validation assays, only weak partial agonism was observed against these receptors, with EC_50_ values greater than 1 µM (**Fig. S9**), 1,000-fold weaker than CB1R. Intriguingly, no agonist activity was seen for the putative cannabinoid receptors GPR55, GPR18, or GPR119. Taken together, **‘4042** appears to be selective for CB1R and CB2R over many other integral membrane receptors.

To minimize locomotor effects in pharmacokinetic exposure experiments, we used a dose of 0.2 mg/kg (**Fig. S10A-B**). At this low dose, **‘4042** was found appreciably in brain and plasma, but not CSF compartments, with higher exposure in brain tissue (AUC_0→inf_ = 3180 ng*min/mL) than plasma (AUC_0→inf_ = 1350 ng*min/mL). The molecule achieved total concentrations in the brain (C_max_ = 16.8 ng/g) and plasma (C_max_ = 5.14 ng/mL or 12 nM) at this dose. A similar pharmacokinetic profile was observed for the positive control CP-55,940 at 0.2 mg/kg, reaching similar maximum concentrations in the brain (C_max_ = 19.2 ng/g versus 16.8 ng/g for **‘4042**), and similar half-lives (T_1/2_ = 127 min versus 114 min for **‘4042**). The main notable difference was seen in the plasma compartment, with a nearly 10-fold increased C_max_ for CP-55,940 compared to **‘4042**. Finally, the concentration of **‘4042** needed to activate the identified off-target receptors even partially is over 10,000-fold higher than the observed gross concentrations, suggesting that activity seen *in vivo* with this ligand reflects on-target engagement (something also consistent with CB1R knockout experiments, below).

#### Anti-allodynia and analgesia

We next tested the efficacy of **‘4042** and its more potent enantiomer **‘1350** *in vivo* in models of acute and chronic pain. We first focused on acute thermal pain. In tail flick, hot plate, and Hargreaves tests of thermal hypersensitivity, **‘4042** and the more potent enantiomer **‘1350** dose-dependently increased tail flick and paw withdrawal latencies. We recorded significant analgesia, namely latencies above baseline, at as little as 0.1 mg/kg dosed intraperitoneally (i.p.) (**Fig. 5A-B, Fig. S11A**). We also recorded increased latencies with the positive control ligand CB1R CP-55,940, but at higher doses (0.5 mg/kg or 0.2 mg/kg doses in the tail flick and Hargreaves tests, respectively.

**Figure 5.**
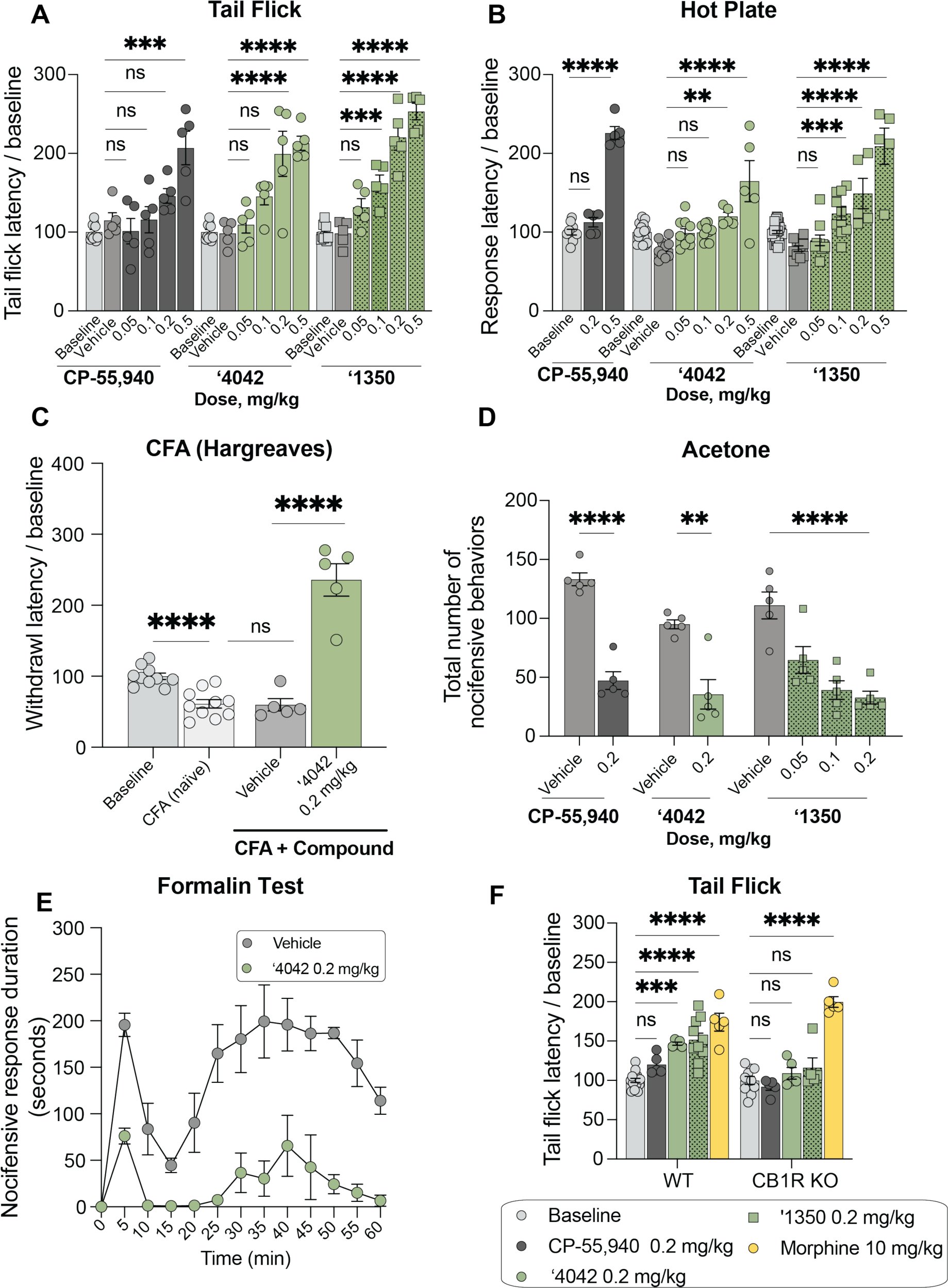
In vivo analgesic profile of ‘4042 and ‘1350. **A.** Dose-response activity in the tail flick assay for CP-55,940 (*n* = 5; one-way ANOVA, *F*(5, 29) = 10.9, *P* < 0.0001), **‘4042** (*n* = 5; one-way ANOVA, *F*(5, 29) = 17.4, *P* < 0.0001) and **‘1350** (*n* = 5; one-way ANOVA, *F*(5, 29) = 48.1, *P* < 0.0001). For all comparisons, asterisks define individual group differences to respective vehicle control using Dunnett’s multiple comparisons post-hoc test correction. **B.** Dose-response activity in the Hot Plate assay for CP-55,940 (*n* = 5; one-way ANOVA, *F*(2, 17) = 148.6, *P* < 0.0001), **‘4042** (*n* = 5-10; one-way ANOVA, *F*(5, 54) = 13.5, *P* < 0.0001) and **‘1350** (*n* = 5-10; one-way ANOVA, *F*(5, 64) = 29.2, *P* < 0.0001). For all comparisons, asterisks define individual group differences to respective vehicle control using Dunnett’s multiple comparisons post-hoc test correction. **C.** Hargreaves test after CFA of **‘4042** (*n* = 5 — 10 per group; two-tailed unpaired *t*-test, **‘4042** versus vehicle: *t*(8) = 7.2, *P* < 0.0001; vehicle versus CFA: *t*(13) = 0.13, *P* = 0.89) after CFA treatment (two-tailed unpaired *t*-test, CFA versus baseline: *t*(18) = 5.2, *P* < 0.0001). **D**. Chemical hyperalgesia test after spared nerve injury. Statistics defined in **Fig. S11** legend. **E.** Nocifensive response duration after formalin treatment (*n* = 5; multiple two-tailed unpaired *t*-tests at each timepoint with the Holm-Šídák post-hoc test correction; all times \**P* < 0.05 – \*\*\*\**P* < 0.0001 except 0 min. and 15 min. (interphase), not significant). **F.** Comparison of the effect of **‘4042**, **‘1350**, CP-55,940, and morphine in wildtype (WT) versus CB1R knockout (KO) mice in the Tail Flick assay (all *n* = 5; two-way ANOVA; genotype x drug treatment interaction: *F*(4, 60) = 6.7, *P* = 0.0002; genotype: *F*(1, 60) = 10.8, *P* = 0.001; drug treatment: *F*(4, 60) = 45.5, *P* < 0.0001; asterisks define individual group differences after Šídák’s multiple comparisons post-hoc test correction).

Next, we assessed the analgesic properties of **‘4042** in a chronic pain model. As illustrated in **Fig. 5C**, 0.2 mg/kg i.p. of **‘4042** was also analgesic in the Complete Freund’s Adjuvant (CFA)-induced inflammatory pain model, increasing paw withdrawal latencies to well-above pre-CFA baseline thresholds. On the other hand, the same 0.2 mg/kg i.p. dose of **‘4042** did not counter the mechanical allodynia that develops in the spared nerve injury (SNI) model of neuropathic pain (**Fig. 5D, Fig. S11B-C**). We did record a modest anti-allodynic effect when dosed intrathecally (i.t.; up to 100 µg/kg; **Fig. S11E-F**), consistent with literature reports of weak effects of other CB1R agonists on mechanical hypersensitivity^44–46^. Furthermore, **‘4042** did not alter the mechanical thresholds of naïve (non-SNI) animals dosed i.p. at 0.2 mg.kg (**Fig. S11D**), a dose that was frankly analgesic in thermal pain assays. Intriguingly, **‘4042**, **‘1350**, and CP-55,940 strongly reduced SNI-induced cold allodynia, another hallmark of neuropathic pain, significantly decreasing the total number of paw withdrawals, a typical acetone-induced nocifensive behavior (**Fig. 5D, S11G**). Finally, in the formalin model of post-operative pain, an i.p. administration of 0.2 mg/kg **‘4042** profoundly decreased the duration of both phase 1 and phase 2 nocifensive behaviors throughout the 60-minute observation period (**Fig. 5E**). We conclude that these new CB1R agonists have strong therapeutic potential across multiple pain modalities in both acute and chronic pain settings.

#### On target activity: CB1R vs CB2R

Consistent with CB1R being the target of **‘4042** and **‘1350** *in vivo*, both total knockout of CB1R as well as pre-treatment with the CB1R selective antagonist AM251 (5.0 mg/kg) completely blocked the analgesic effect of **‘4042** and **‘1350**, but not of morphine, in the tail flick assay (**Fig. 5F, S11H**). In contrast, neither CB2R knockout nor co-treatment with the CB2R-selective antagonist SR-144528 (1.0 mg/kg) decreased the analgesic effects of **‘4042** in the tail flick or Hargreaves assays (**Fig. S11I-K**). We conclude that the anti-allodynic (cold/SNI), anti-hyperalgesic (CFA) and analgesic (thermal, acute) effects of **‘4042** and **‘1350** are CB1R, and not CB2R, dependent.

#### Cannabinoid tetrad of behaviors

The cannabinoid “tetrad” of behaviors is widely used to assess CNS engagement of cannabinoid receptors by new agonists^11^. This suite of tests measures the four hallmarks of CB1R agonism, namely analgesia and three common cannabinoid side-effects—hypothermia, catalepsy, and hypolocomotion. Given the new chemotypes discovered here, we also examined our leads **‘4042** and **‘1350**, in this panel of potential side-effects.

#### Reduced sedation at analgesic doses

Hypolocomotion, one of the four features of the tetrad, is a commonly assessed proxy for the sedative side-effect of cannabinoids.Sedation is not only an important clinical adverse side effect of cannabinoids, but it also confounds preclinical reflex tests of analgesia, where unimpeded movement of a limb is the endpoint. Intriguingly, although mice treated with ‘**4042** appeared less active than those treated with vehicle, ‘**4042**-injected mice were not sedated at analgesic doses (**Fig. 6A-B**). Not only did the mice promptly move when slightly provoked (touched, or their housing cylinders slightly disturbed), but in two quantitative and widely-used assays of hypolocomotion and sedation, the open field and rotarod tests, we found no significant differences between ‘**4042**- and vehicle-treated animals at analgesic doses (**Fig. 6A, D**). Higher doses did tend to decrease overall locomotor activity, but only at the highest dose (1.0 mg/kg) did we record some motor deficits in the rotarod test (**Fig. 6B, E**). We do note that the more potent pure enantiomer, **‘1350,** did show locomotor deficits at lower doses: 0.2 mg/kg in the open field test (**Fig. 6A**) and 0.5 mg/kg in the rotarod test (**Fig. 6B**), these are balanced by **‘1350**’s increased potency at CB1R (**Fig. 4**) and its stronger analgesia at equivalent doses to **‘4042** (**Fig. 5A-B,D**). In contrast, all analgesic doses tested with the positive control CP-55,940 caused motor impairment in both the rotarod and open field tests (**Fig. 6A-B, D-E**), suggesting that the analgesia produced by CP-55,940 is confounded by its sedative effect (**Fig. 5A,D, S11A**).

**Figure 6.**
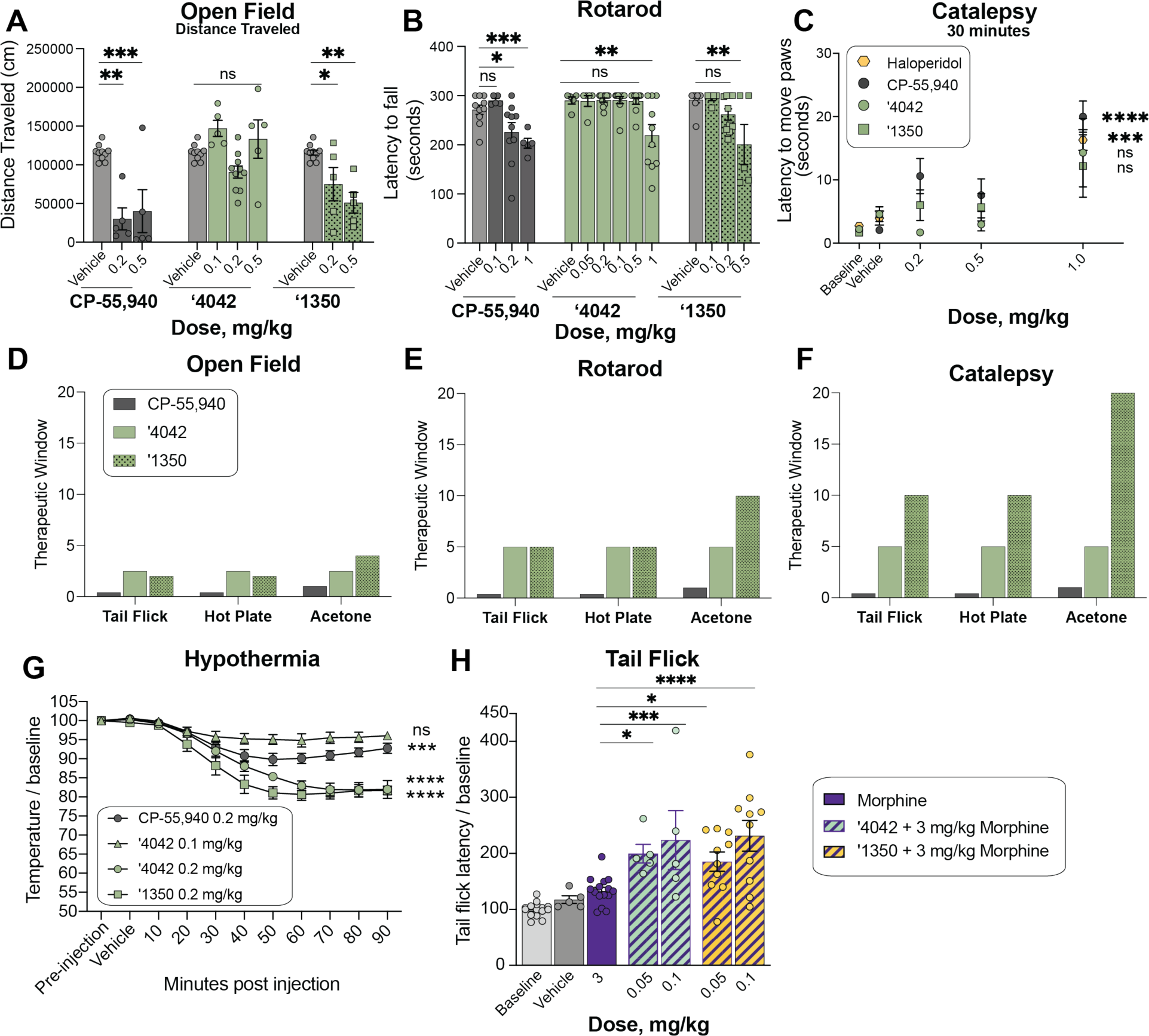
In vivo side-effect and cotreatment profile of ‘4042 and ‘1350. **A.** Dose-response of **‘1350** (all *n* = 5; one-way ANOVA, *F*(2, 17) = 9.5, *P* = 0.002), **‘4042** (0.1 and 0.5 mg/kg, *n* = 5; 0.2 mg/kg *n* = 10; one-way ANOVA, *F*(3, 26) = 5.3, *P* = 0.006) and CP-55,940 (all *n* = 5; one-way ANOVA, *F*(2, 17) = 13.7, *P* < 0.001) in the open-field test of hypolocomotion. Asterisks define individual group differences to vehicle control after Dunnett’s multiple comparisons post-hoc test correction. **B.** Rotarod test of sedation comparison of CP-55,940 (all *n* = 5 except 0.2 mg/kg *n* = 10; one-way ANOVA, *F*(3, 26) = 5.7, *P* = 0.04) to **‘4042** (all *n* = 10 except 0.05 mg/kg *n* = 5; one-way ANOVA, *F*(5, 44) = 6.2, *P* < 0.001) and **‘1350** (all *n* = 5 except 0.2 mg/kg *n* = 10; one-way ANOVA, *F*(3, 26) = 5.7, *P* = 0.004); asterisks define individual group differences to respective vehicle control after Dunnett’s multiple comparisons post-hoc test correction. **C.** Mesh grip test of catalepsy at 30 minutes post-dose. Comparison of CP-55,940 (*n* = 5-10; one-way ANOVA, *F*(3, 26) = 10.7, *P* < 0.0001), haloperidol (*n* = 5; two-tailed unpaired *t*-test, *t*(8) =6.2, *P* < 0.001), **‘4042** (*n* = 5; one-way ANOVA, *F*(3, 16) = 4.1, *P* = 0.02) and **‘1350** (*n* = 5-10; one-way ANOVA, *F*(3, 26) = 1.02, *P* = 0.4). Asterisks define differences between 1 mg/kg dose for each compound and respective vehicle control. Data at 1 hr timepoint are in **Fig. S11**. **D-F.** Therapeutic windows for each analgesia test versus hypolocomotion (**D.**, open field), sedation (**E.**, rotarod), and catalepsy (**F.**) side-effects. Therapeutic window was calculated as the ratio of the minimum dose of side-effect onset or maximum tested side-effect dose if no doses were significant to the minimum dose of analgesia onset. **G.** Body temperatures of mice treated with CP-55,940 (*n* = 5; one-way ANOVA, *F*(10, 44) = 13.3, *P* < 0.0001), **‘4042** (0.1 mg/kg; *n* = 5; one-way ANOVA, *F*(10, 44) = 3.5, *P* = 0.002; 0.2 mg/kg; *n* = 5; one-way ANOVA, *F*(10, 44) = 32.2, *P* < 0.0001) and **‘1350** (*n* = 3; one-way ANOVA, *F*(10, 22) = 27.3, *P* < 0.0001). Asterisks define differences between each group at 90 min. post-dose and their respective vehicle control. mg/kg dose for each compound and respective vehicle control. **H.** Cotreatment of subthreshold dose of morphine with **‘4042** (one-way ANOVA, *F*(4, 40) = 11.0, *P* < 0.0001) and **‘1350** (one-way ANOVA, *F*(4, 50) = 14.7, *P* < 0.0001) in the tail flick test. Asterisks define cotreatment differences to morphine alone (3 mg/kg) using Dunnett’s multiple comparisons post-hoc test correction. For all statistical tests: ns, not significant, \**P* < 0.05, \*\**P* < 0.01, \*\*\**P* < 0.001, \*\*\*\**P* < 0.0001.

#### Reduced catalepsy at analgesic doses

To determine whether **‘4042** or **‘1350** induce catalepsy, we measured the latency of compound-injected mice to move all four paws when placed on a vertical wire mesh. As expected, mice injected with the non-cannabinoid positive control haloperidol induced dramatic catalepsy (**Fig. 6C, S11L**). Conversely, and consistent with their decreased locomotor effects, analgesic doses (0.2 or 0.5 mg/kg) of **‘4042** and **‘1350** did not induce cataleptic behavior, 30 minutes or 1-hour post-injection. In fact, we recorded catalepsy only at doses that also induced motor deficits (i.e, > 1.0 mg/kg), which may have contributed to the increased latency to move the paws. In contrast, CP-55,940 exhibited catalepsy starting at 0.2 mg/kg (**Fig. 6C**), consistent with the effects seen in the open field and rotarod tests (**Fig. 6A-B**); here too, there was no window between analgesia and catalepsy for this widely-used cannabinoid probe (**Fig. 6F**).

#### ‘4042 and ‘1350 induce hypothermia

Finally, we examined the effect of the new CB1R agonists on hypothermia. Here, we measured body temperature of mice implanted with telemetric probes continuously for 150 minutes. All three of CP-55,940, **‘4042**, and **‘1350** induced hypothermia compared to baseline and vehicle (**Fig. 6G**).

Overall **‘1350** and **‘4042** had reduced adverse reactions at analgesic doses versus the classic cannabinoid CP-55,940. For the classic adverse “tetrad” behaviors, CP-55,940 induced meaningful catalepsy and sedation at the same concentrations where it conferred anti-allodynia and analgesia; for this widely used cannabinoid it was impossible to deconvolute effects on pain from the adverse effects. This is as expected and is why the “tetrad” is considered characteristic of active cannabinoids. Conversely, depending on the nociceptive behavior, **‘1350** and **‘4042** had up to a twenty-fold concentration window between anti-allodynia or analgesia versus catalepsy and sedation, and typically a five- to ten-fold concentration window (**Fig. 6D-F**). This is most noticeable in the acetone test for cold pain perception, where **‘1350** demonstrated significant anti-allodynia at 0.05 mg/kg but only began to show increased latency to move paws, suggestive of catalepsy, at 1 mg/kg doses. In heat-based nociception, both in the tail-flick, which is reflex-based, and hot-plate, which is more affective, **‘1350** had at least a ten-fold window between anti-allodynia (significant at 0.1 mg/kg) and catalepsy (1 mg/kg highest tested dose) (**Fig. 6F**). In other behaviors the window dropped, for instance between heat-based response and sedation as measured by the rotarod, it was only five-fold (**Fig. 6E**). However, in almost every behavior there was a meaningful window between nociception versus catalepsy and sedation, which is rare among classic cannabinoids such as CP-55,940.

#### Pretreatment with ‘4042 or ‘1350 increases the analgesic effect of morphine

As cannabinoids have been shown to potentiate morphine analgesia, we investigated whether co-treatment of ‘**4042** or **‘1350** with morphine has better pain-relieving properties than morphine alone. Here, we combined low doses (0.05 and 0.1 mg/kg) of **‘4042** or **‘1350** with morphine (3.0 mg/kg, i.p.) and tested the analgesic efficacy of the combination vs. morphine alone in the tail flick assay. Mice co-injected with any combination of morphine and ‘**4042** or **‘1350** exhibited significantly longer tail flick latencies than did mice injected with morphine alone (**Fig. 6H**). This result suggests that these combinations have at least an additive analgesic effect when combined, consistent with previous studies on circuitry^47^ and CB1/2R ligand polypharmacy with morphine^47–49^.

#### The new CB1R agonist is not rewarding

A major limiting factor in an analgesic’s clinical utility, particularly opioids, is misuse potential because of rewarding properties. To determine whether ‘**4042** exhibits comparable liabilities, we turned to the conditioned place preference (CPP) test in which mice learn to associate one chamber of the apparatus with a compound. If mice show a preference for the drug-paired chamber, then the compound is considered to be intrinsically rewarding. As expected, mice injected with morphine significantly increased their preference for the chamber associated in which they received the drug versus its vehicle-associated chamber (**Fig. S11M**). In contrast, mice injected with ‘**4042** spent comparable time in the ‘**4042**-paired or vehicle-paired chambers, indicating that ‘**4042** does not induce preference at these doses. Similarly, we found that mice injected with the cannabinoid CP-55,940 did not spend more time in the drug-paired chamber; in fact, mice spent significantly more time in the vehicle-paired chamber, suggesting that CP-55,940 may actually induce some aversion, something not seen with **‘4042** but consistent with previous studies using a similar dose range^50,51^.

## DISCUSSION

From a vast library of virtual molecules, structure-based discovery has led to new agonists that not only potently activate CB1R but are also strongly analgesic without key liabilities of classic cannabinoids. Three observations merit emphasis. **First**, from a tangible library of previously unsynthesized, new to the planet molecules, structure-based docking found new chemotypes for the CB1 receptor, physically distinct from previously known ligands. Using structural complementarity, and the wide range of analogs afforded by the new libraries, we optimized these new ligands, leading to a 1.9 nM K_i_ full agonist of the CB1R. **Second**, the pose adopted by active enantiomer of **‘4042** (**‘1350**) in a cryo-EM structure of its complex with CB1R-G_i_ superposed closely on the docking prediction, explaining its SAR at atomic resolution and supporting future optimization. **Third**, while the racemic agonist **’4042** is strongly anti-allodynic and analgesic across a panel of nociception behavioral assays, it spares several of the characteristic adverse drug reactions of most cannabinoid analgesics, with typically a 5-10-fold window between analgesia and both sedation and catalepsy. Interestingly, **‘1350** exhibited a similar therapeutic window as **’4042** but with a shift towards lower doses; i.e **‘1350** exhibited stronger analgesic effects across multiple tests but also induced side effects at lower doses. These traits are unusual for cannabinoids, where sedation often closely tracks with analgesia and where catalepsy is among the “tetrad” of side-effects characteristic of cannabinoid agonists. Encouragingly, administration of morphine with low doses of **‘4042** or **‘1350** show improved analgesia, suggesting that the combination of low doses of opioids and cannabinoids retains significant analgesia but potentially with a lower side effect profile, therefore expanding the therapeutic window of each compound on its own.

### Limitations of this Study

Several caveats bear mentioning. The mechanistic bases for the disentanglement of sedation and catalepsy from analgesia remains uncertain. Often, clear differences in functional or subtype selectivity support phenotypic differences of different ligands^26,27,41,52^. Here, functional differences between **‘4042** and **‘1350,** which show two reduced characteristic “tetrad” behaviors, and CP-55,940, which does not, were modest, with only notable differences shown at CB1R for recruitment of G_13_. The functional importance of G_13_ in the *in vivo* models is not understood but could be explored in the future. Pronounced differences were, however, seen in the functional effects between the CB1R and CB2R subtypes. Though it is possible that the described CB2R partial agonism could be a feature that separates **‘4042** from CP-55,940 and other cannabinoids, studies in cannabinoid receptor knockout animals suggest that catalepsy and sedation are completely ablated in CB1R, but not CB2R mice^53^. Additionally, in our hands using CB2R knockout mice, at minimum the analgesic effects are not due to engagement of CB2R receptors. The contribution of other off-targets, such as antagonism of GPR55 or engagement of TRPV1, may merit further exploration. Still, without a definitive molecular mechanism at this time, we can for now only lay the ability to disentangle analgesic efficacy from “tetrad” adverse reactions at the door of the new chemotypes explored^54–56^.

Despite these caveats, the main observations of this study are clear. Docking a large library of virtual molecules against CB1R revealed new agonist chemotypes, the most promising of which was optimized to the potent full-agonist **‘4042.** A cryo-EM structure of the ***R-‘*4042**-CB1-G_i1_ complex confirmed its docking-predicted pose. The new agonist is strongly analgesic, and unlike most cannabinoids generally has a 5-10-fold therapeutic window over sedation and catalepsy. We suspect that newer chemotypes still remain to be discovered, and that these might further separate the dose-limiting side-effect aspects of the cannabinoid tetrad while maintaining analgesic potency, supporting the development of new cannabinoid medicines to treat pain.

## Acknowledgements

This work is supported by DARPA grant HR0011-19-2-0020 (B.K.S., A.I.B., & J.J.I.), US NIH grant R35GM122481 (B.K.S.), US NIH grant R01GM71896 (J.J.I.), US R35NS097306 (A.I.B.), Open Philanthropy (A.I.B.), the Facial Pain Research Foundation (A.I.B.), and US NIH grant P01DA009158 (A.M.). We thank C. Webb for help with the initial CB1R docking screen. We thank B. Ahanou for help analyzing the open field data. We thank M. M. Rachman for editing the manuscript. We gratefully acknowledge OpenEye Software for Omega and related tools and Schrödinger, Inc. for the Maestro package. Select receptor binding profiles and agonist functional data was generously provided by the National Institute of Mental Health’s Psychoactive Drug Screening Program, Contract # HHSN-271-2018-00023-C, directed by B. Roth.

## Author contributions

T.A.T. and R.M.S. conducted the docking screens with input from B.K.S. Ligand optimization was performed by T.A.T. and C.I.-T. with input from B.K.S., and A.M.. N.G.T., S.G., F.T., and Y.L. performed binding or functional assays with input from T.A.T., C.I.-T., and A.M.. K.K. prepared the CB1-G_i_ complex, collected cryo-EM data with help from E.S.O., and modelled the structure with help from F.L.. K.K. collected bimane data with help from YS.. T.A.T. and J.M.B. did the drug formulations for in vivo experiments. J.M.B. performed and analyzed the *in vivo* pharmacology experiments assisted by T.A.T., V.C., S.R.R., K.B., and J.B., supervised and coanalyzed by A.I.B.. E.S.O processed data and obtained the cryo-EM map. Y.S. performed the GTP-turnover assays with help from K.K. and E.S.O.. H.K. and C.N. tested select compounds in the panel of G-protein and β-arrestin subtypes with supervision from M.S. and L.S.. C.I.-T. and T.C.H. synthesized bespoke compounds with supervision from A.M.. I.G. performed the colloidal aggregation screens. Y.S.M. and D.S.R. supervised compound synthesis of Enamine compounds purchased from the ZINC15 database and 12 billion catalog. J.J.I. built the ZINC15 ultra-large libraries. B.K.S., A.I.B., A.M., and K.K. supervised the project. T.A.T. wrote the paper with input from all other authors, and primary editing from B.K.S.

## Supplemental Data Figures

**Figure S1.**
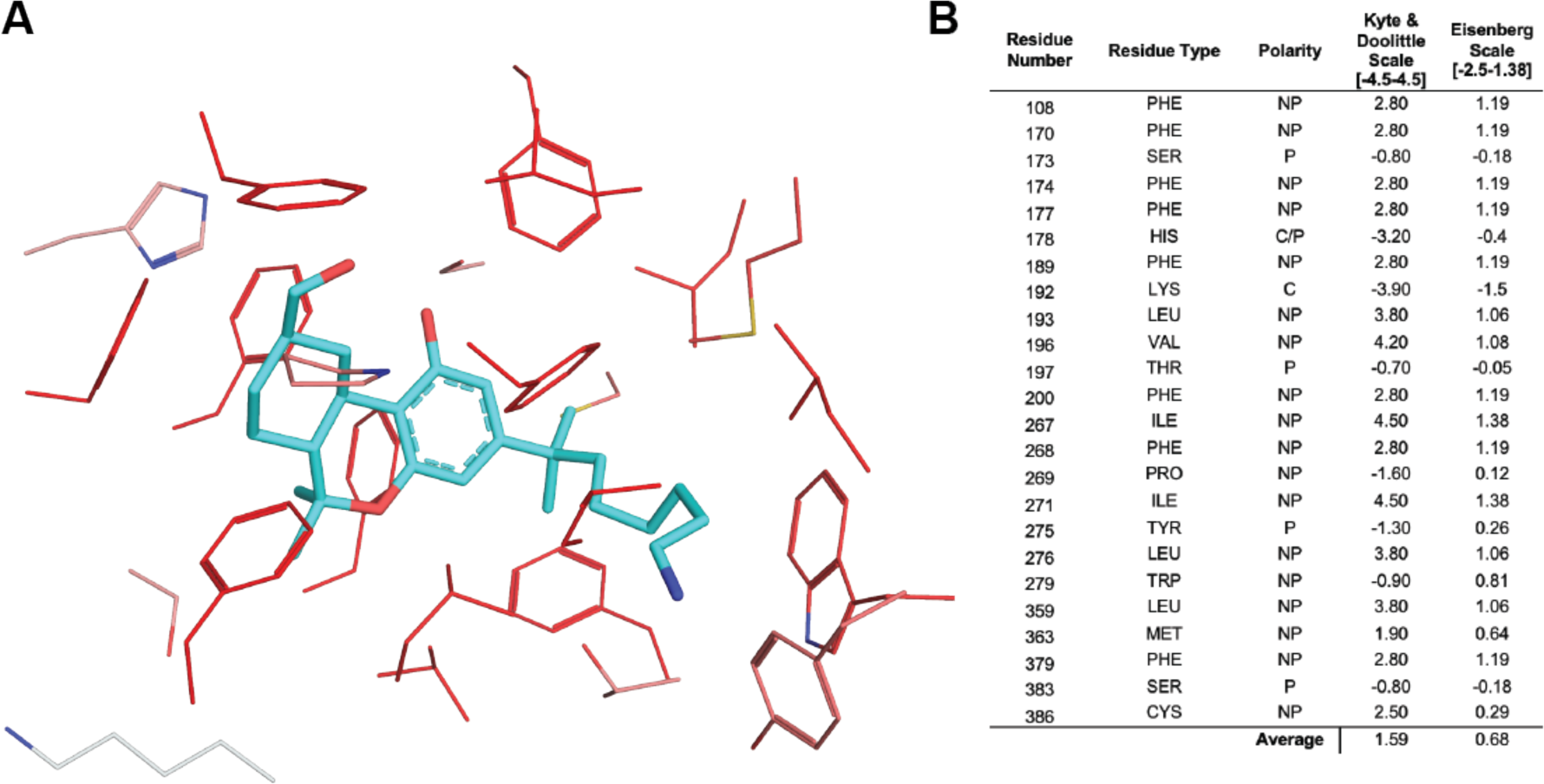
Hydrophobicity calculations for the hCB1R orthosteric pocket based on PDB: 5XR8. Residues within 5 Å of AM841 are considered. **A.** Depiction of the hCB1 orthosteric pocket, colored by the Eisenberg Scale, where darker red colors indicate more hydrophobic residues and lighter red or gray colors indicate less hydrophobic residues. **B.** A table of the residues within 5 Å of AM841, with their polarity class, and two hydrophobicity scores indicated.

**Figure S2.**
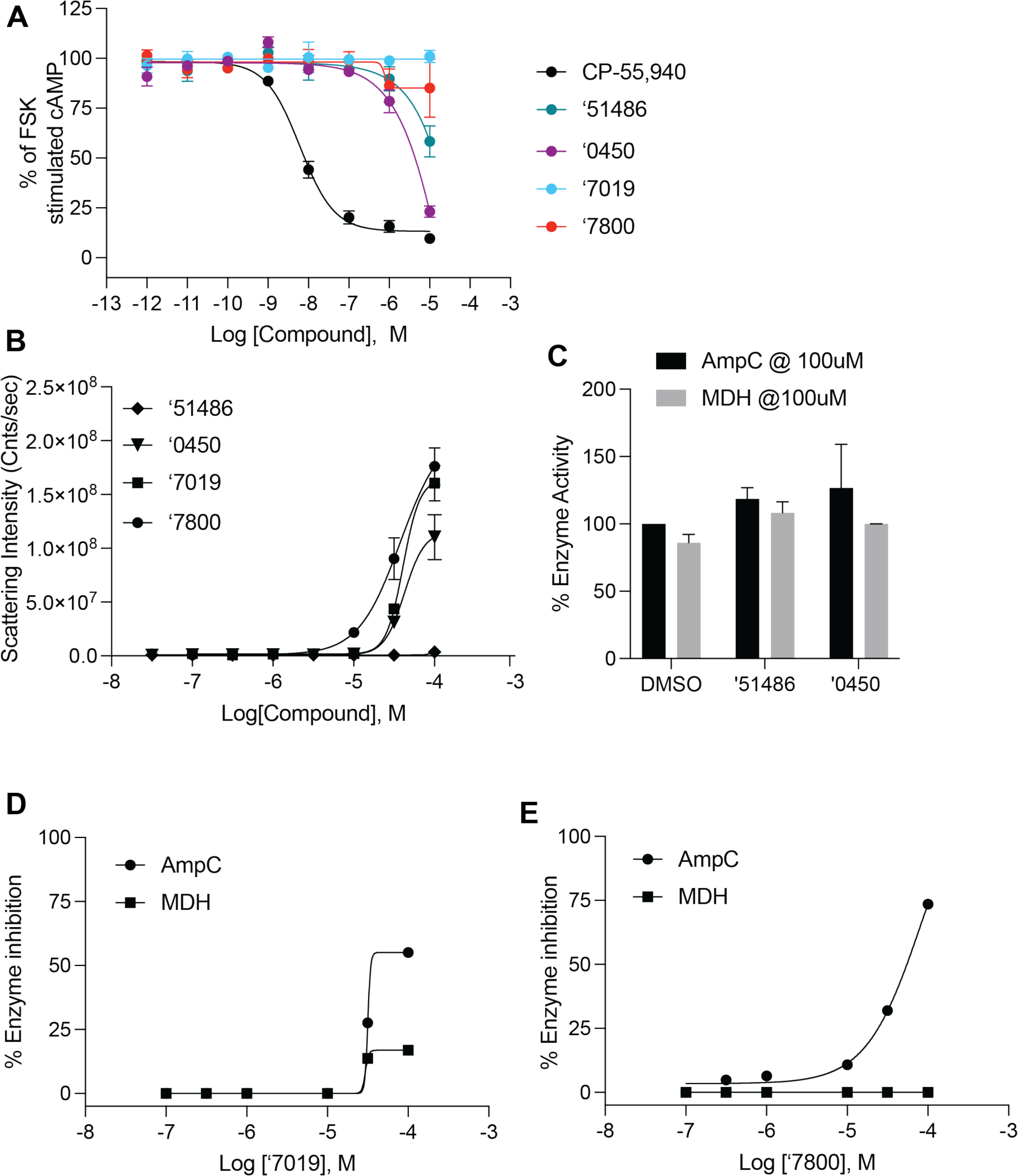
Functional measurements for a subset of screening hits. **A.** Functional cAMP inhibition at hCB1R by the four most potent docking hits. **B.** Scattering intensity in dynamic light scattering experiments of colloidal aggregation. **C.** Inhibition of the off-target enzymes MDH and AmpC Beta-lactamase at 100 uM. **D.** and **E.** Single-point inhibition of the off-target enzymes MDH and AmpC Beta-lactamase by **‘7019 (D.)** and **’7800 (E.).** All data represent mean ± SEM of three independent experiments in triplicate except **B.** which represents one independent experiment in triplicate.

**Figure S3.**
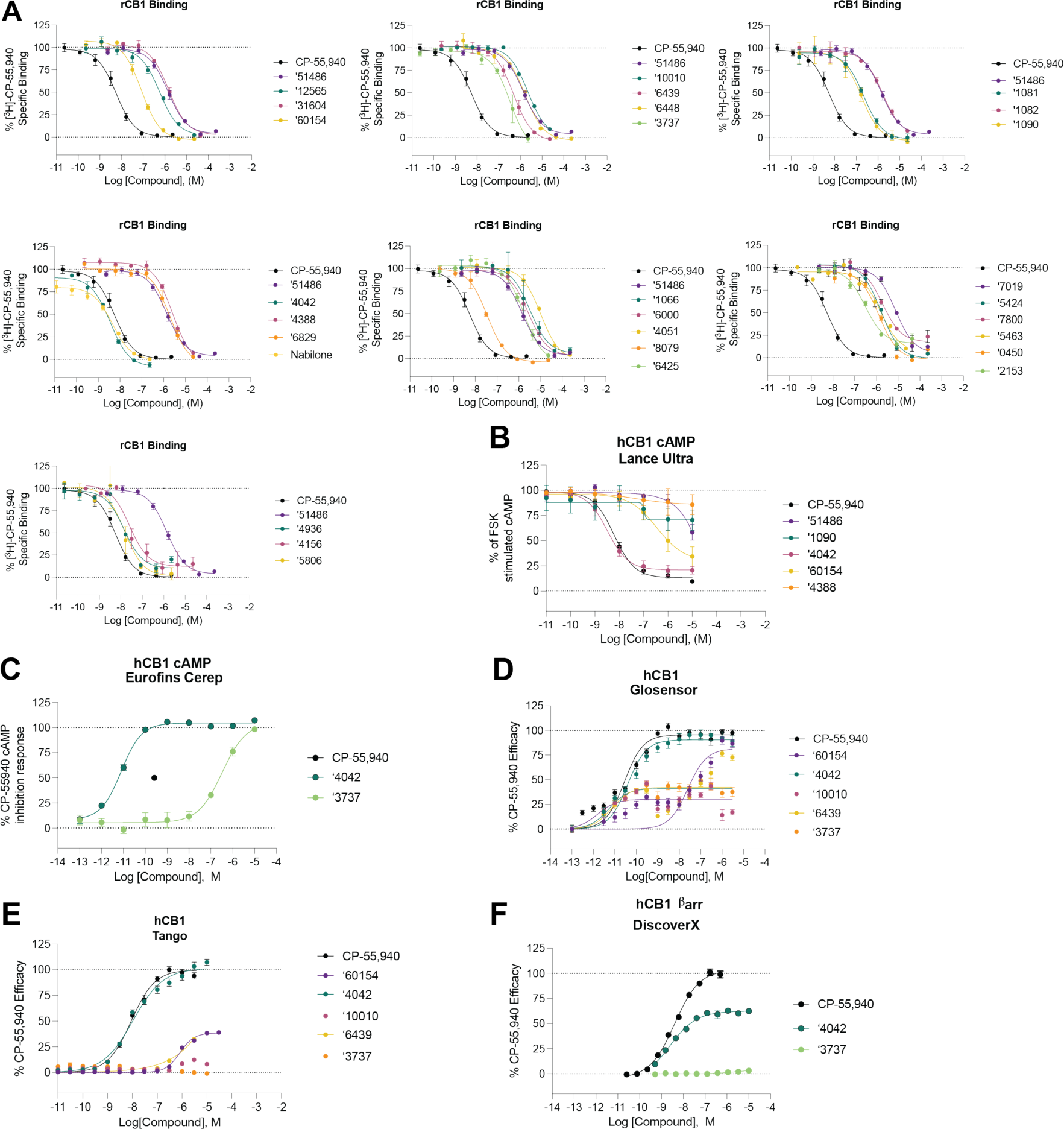
hCB1 binding and functional data for analogs. **A.** Competition binding data for primary hits and a subset of their analogs at hCB1. **B-D.** Functional cAMP inhibition for a subset of analogs at hCB1 across three separate assays. **E-F.** Functional ß_arr_ recruitment for a subset of analogs. All data represent mean ± SEM of at least 2 independent experiments in triplicate except **C.** and **F.** which represent one independent experiment in triplicate. Best fit values can be found in **Supplementary Table 2**.

**Figure S4.**
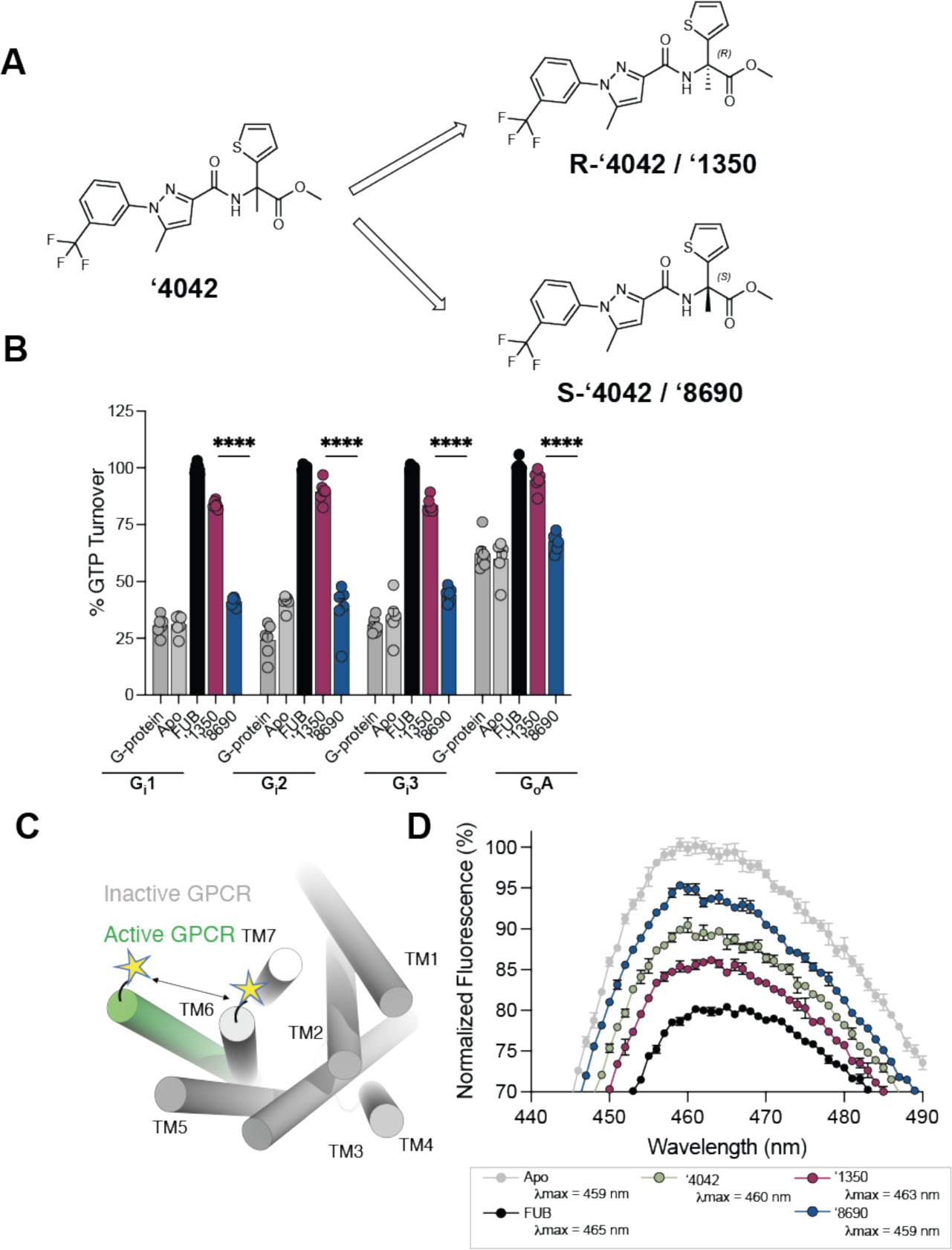
Additional pharmacological characterization of **‘4042** and its enantiomers. **A.** Chiral column purification led to the separation of two independent enantiomers, **’1350** and **‘8690**. ‘**1350** was determined to be ***R*-’4042** from the Cryo-EM structure. **B.** GTPase Glo assay characterizing GTP turnover of G-proteins G_i1-3/o_. **C.** Schematic of the environmentally sensitive fluorophore Monobromobimane (Bimane) which when site-specifically labeled (e.g. on TM6) acts as a conformational reporter. **D**. Compared to the apo (grey), the spectrum of full agonist MDMB-fubinaca (Fub)-bound CB1 (black) shows a decrease in intensity and a blue-shift in λ_max_ (Apo 459 nm to Fub 465 nm). The bimane spectrum of **‘8690** (λ_max_ 459 nm, blue) is more similar to apo and the spectrum of **‘1350** (λ_max_ 463 nm, magenta) is closer to that of Fub. The spectrum of the racemate, **‘4042** (green) is between **’1350** (***R-*‘4042**) and **‘8690** (***S-*‘4042**). All data represent mean ± SEM of three independent experiments in triplicate.

**Figure S5.**
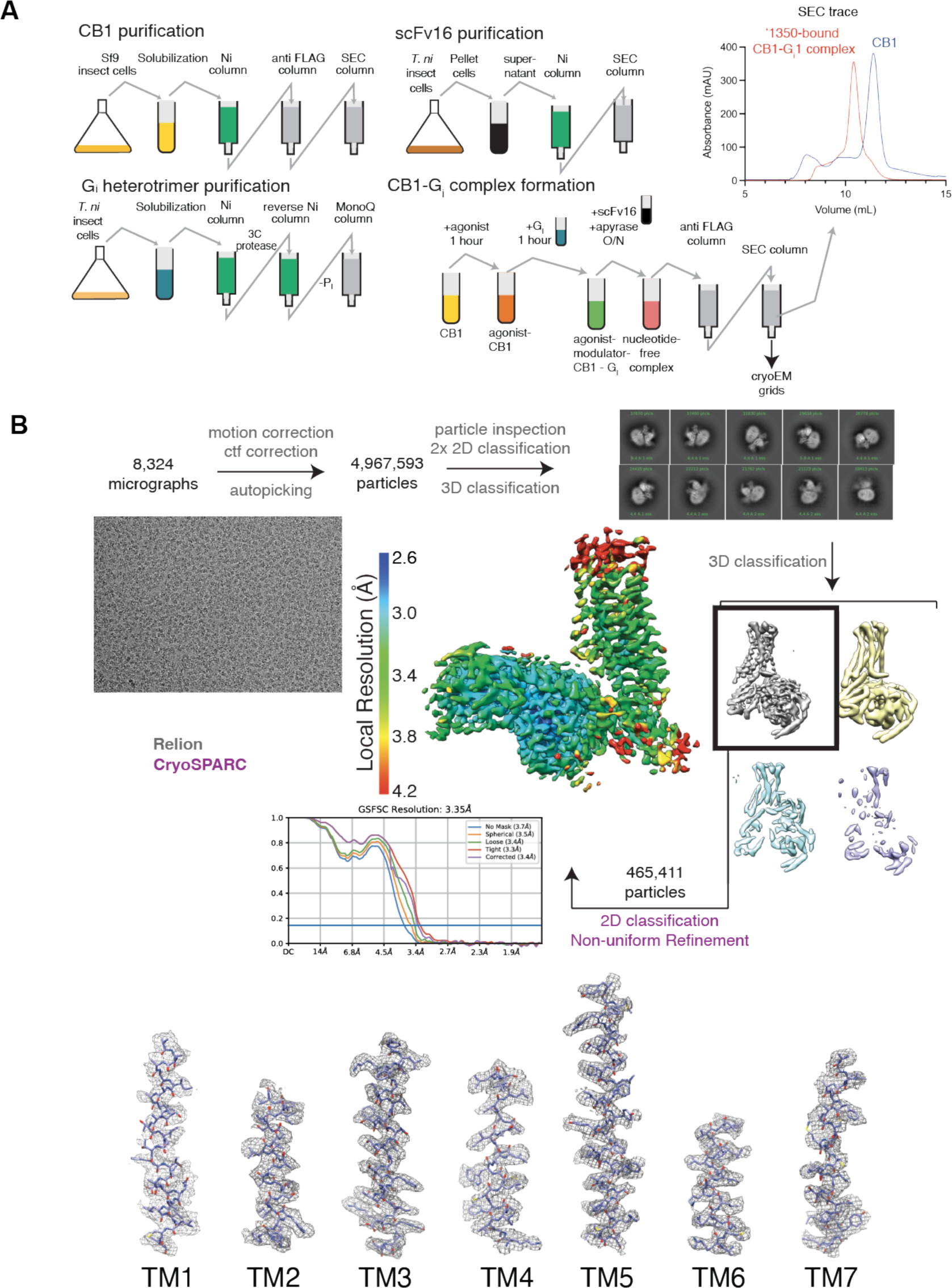
Cryo-EM sample preparation and data processing. **A.** Purification of hCB1, scFv16, the G_i_ heterotrimer, and complex formation protocols. **B.** Cryo-EM data processing flow chart of CB1, including particle selection, classifications, and density map reconstruction. Details can be found in **Supplementary Table 3**.

**Figure S6.**
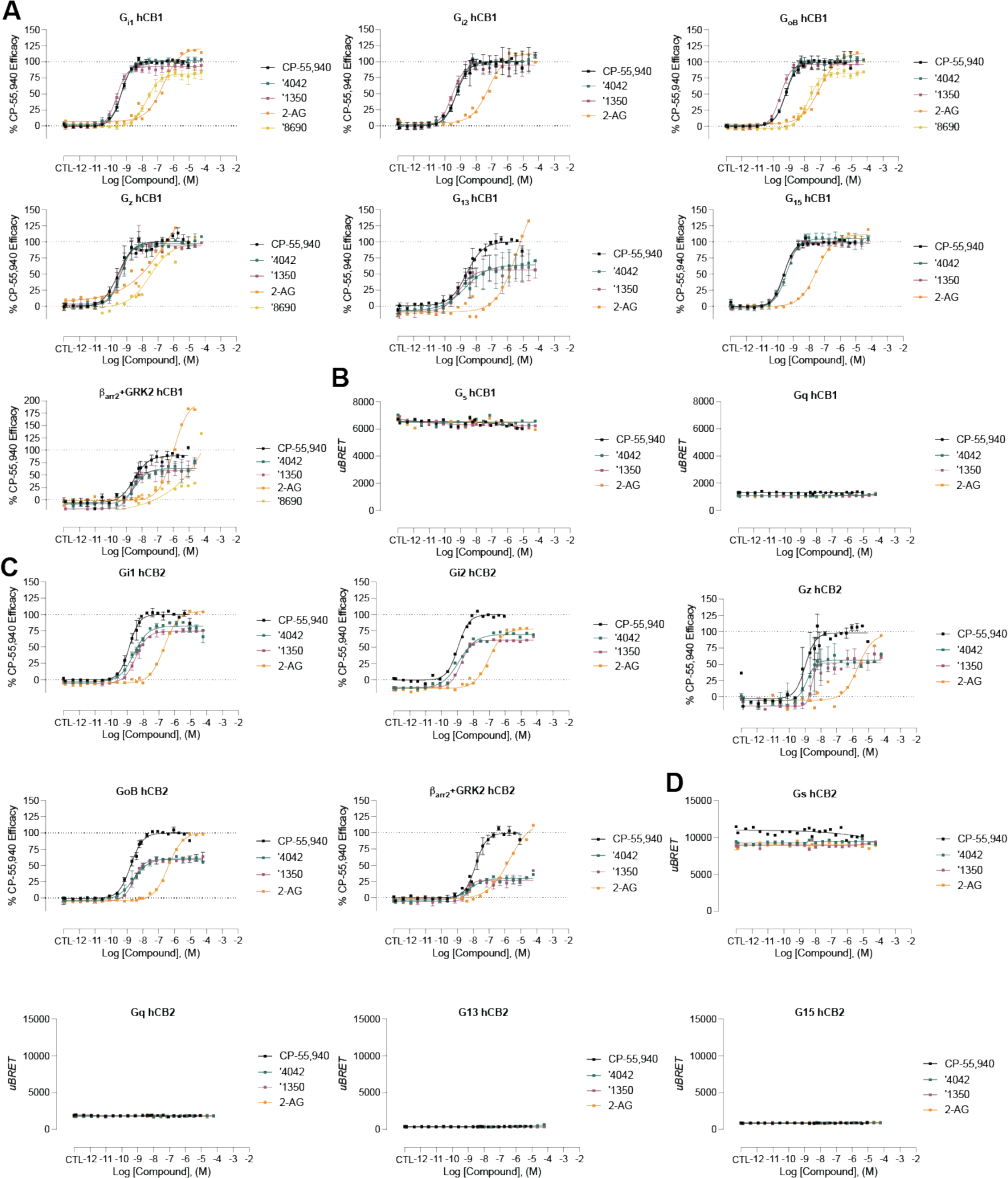
hCB1/2 functional data for select analogs in the bioSens-All^®^ platform. **A.** Normalized activity for select analogs versus a panel of sensors in hCB1-expressing cells. **B.** Raw BRET activity for select analogs versus G_s_ and G_q_ in hCB1-expressing cells. **C.** Normalized activity for select analogs versus a panel of sensors in hCB2-expressing cells. **D.** Raw BRET activity for select analogs versus G_s_, G_q_, G_12_, and G_15_ in hCB2-expressing cells. Best fit values can be found in **Supplementary Tables 5 & 8**.

**Figure S7.**
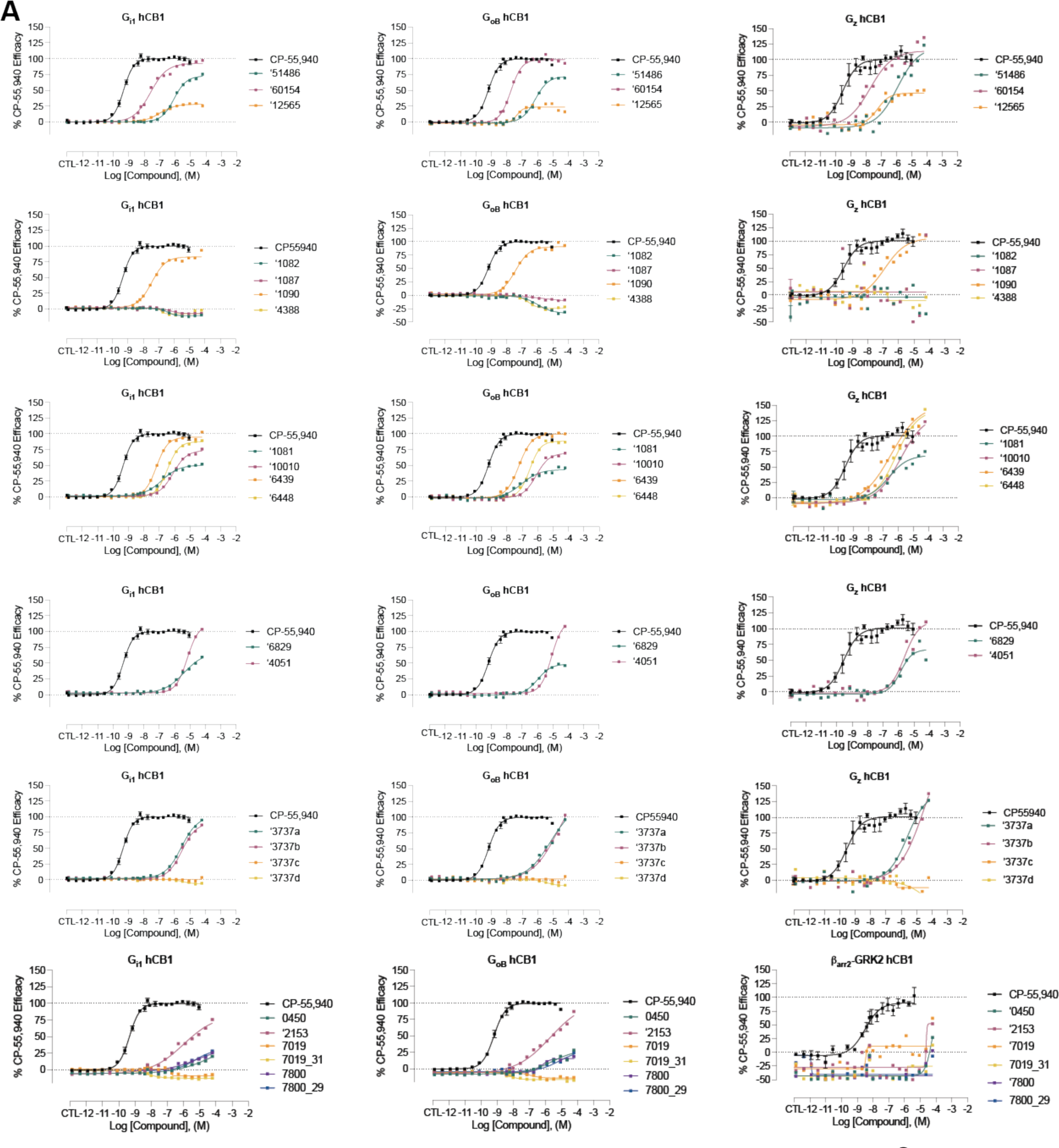
hCB1 functional data for select analogs in the bioSens-All^®^ platform. **A.** Normalized activity for select analogs versus a panel of sensors in hCB1-expressing cells. Best fit values can be found in **Supplementary Table 4**.

**Figure S8.**
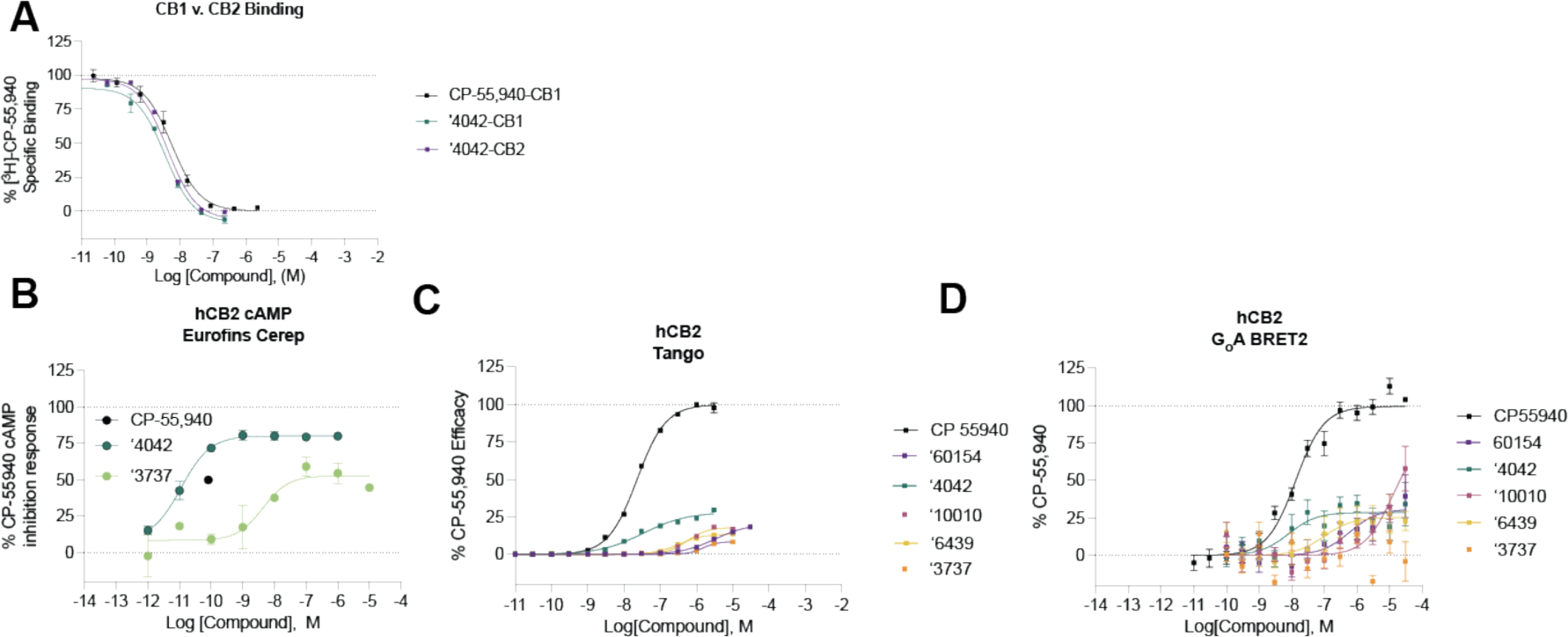
CB2R binding and functional data for select analogs. **A.** Competition binding data shows that **‘4042** is modestly more potent at CB1R than CB2R (rCB1 pKi = 8.7 (95% CI 8.60 – 8.86), hCB2 pKi = 8.6 (95% CI 8.55 – 8.77); *t*(4) =6.5, *p* = 0.003). **B-D.** Functional cAMP inhibition for a subset of analogs at hCB2 across three separate assays. All data represent mean ± SEM of three independent experiments in triplicate except **B.** which represents one independent experiment in triplicate. Best fit values can be found in **Supplementary Table 7**.

**Figure S9.**
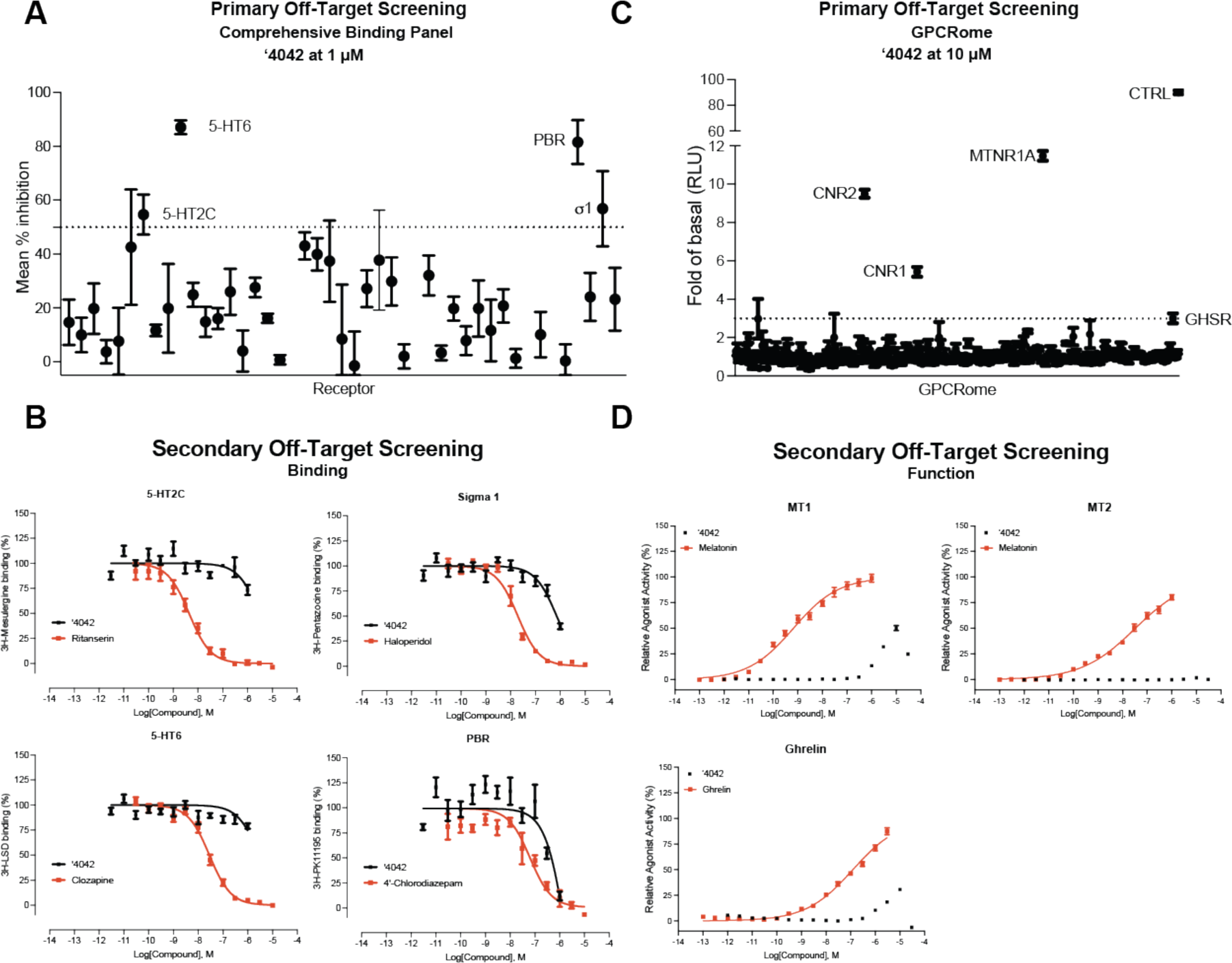
Off-target profiling of **‘4042**. **A.** Comprehensive binding data against a panel of 45 common GPCR and non-GPCR drug targets. **B.** Follow-up dose response binding experiments for targets with > 50% inhibition in the single-point experiments. **C.** TANGO screens against a panel of 320 GPCRs for **’4042**. **D.** Follow-up dose response functional experiments for targets with > 3-fold activation in the single-point experiments. Data in **A.**, **C.,** and **D.** represent mean ± SEM of 3 independent experiments in triplicate. Data in **B.** represent mean ± SEM of 2 independent experiments in triplicate except 5-HT6 which is 3 independent experiments in triplicate.

**Figure S10.**
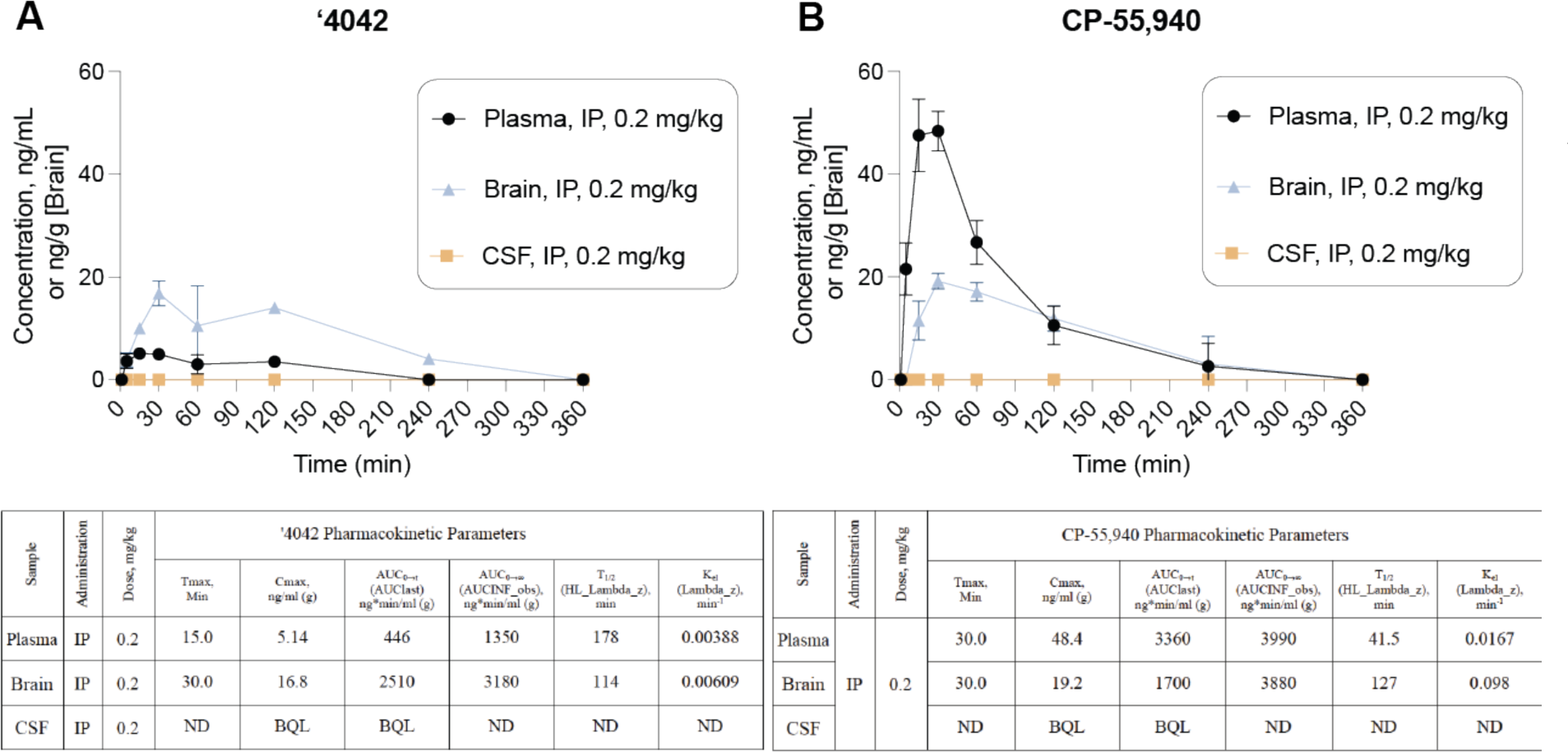
Pharmacokinetic profiles of **‘4042** compared to CP-55.940. Pharmacokinetic profile of **‘4042** (**A**.) and CP-55,940 (**B**.) after a single 0.2 mg/kg dose in brain, CSF, and plasma compartments. Data represent mean ± SEM of 3 animals per timepoint.

**Figure S11.**
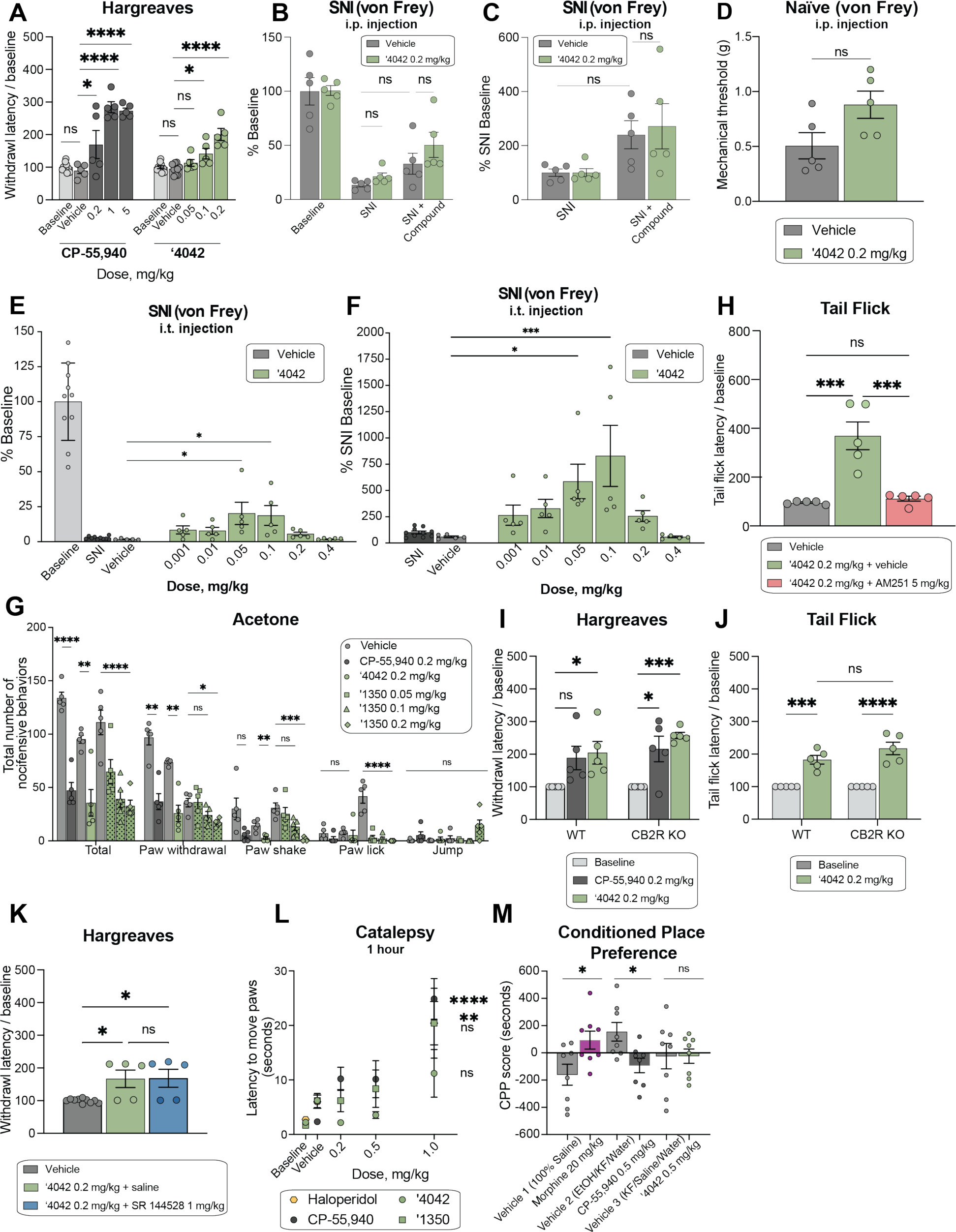
Additional analgesic and side-effect profiles of **‘4042** and **‘1350. A.** Dose-response activity in the Hargreaves assay for **‘4042** (*n* = 5; one-way ANOVA, *F*(3, 21) = 16.3, *P* < 0.0001) and CP-55,940 (*n* = 5; one-way ANOVA, *F*(4, 25) = 26.2, *P* < 0.0001). Asterisks define individual group differences to respective vehicle control using Dunnett’s multiple comparisons post-hoc test correction. **B.** Effect of **‘4042** (i.p.) in neuropathic pain model in mice after SNI with mechanical allodynia (*n* = 5; two-way ANOVA; SNI x drug treatment interaction: *F*(2, 24) = 0.5, *P* > 0.05; SNI: *F*(2, 24) = 51.8, *P* < 0.0001; drug treatment: *F*(1, 24) = 1.6, *P* > 0.05; asterisks define individual group differences to vehicle control after Tukey’s multiple comparisons post-hoc test correction). Data presented are normalized to pre-SNI baseline measurements. **C.** Effect of **‘4042** (i.p.) in neuropathic pain model in mice after SNI with mechanical allodynia (*n* = 5; two-way ANOVA; SNI x drug treatment interaction: *F*(1, 16) = 0.1, *P* > 0.05; SNI: *F*(1, 16) = 9.6, *P* = 0.007; drug treatment: *F*(1, 16) = 0.1, *P* > 0.05; asterisks define individual group differences to vehicle control after Tukey’s multiple comparisons post-hoc test correction). Data presented are normalized to post-SNI baseline measurements. **D.** Effect of **‘4042** (i.p.) in naïve (non-SNI) mice in the mechanical assay (all *n* = 5; two-tailed unpaired t-test, *t*(8) = 2.17, *P* > 0.05). **E.** Effect of **‘4042** (i.t.) in neuropathic pain model in mice after SNI with mechanical allodynia (*n* = 5; one-way ANOVA, *F*(6, 28) = 4.2, P = 0.004; asterisks define individual group differences to vehicle control after Dunnett’s multiple comparisons post-hoc test correction). Data presented are normalized to pre-SNI baseline measurements. **F.** Effect of **‘4042** (i.t.) in neuropathic pain model in mice after SNI with mechanical allodynia (*n* = 5; one-way ANOVA, *F*(7, 32) = 3.8, *P* = 0.004; asterisks define individual group differences to vehicle control after Dunnett’s multiple comparisons post-hoc test correction). Data presented are normalized to post-SNI baseline measurements. **G.** Chemical hyperalgesia test after spared nerve injury (all *n* = 5; **‘4042** vs. vehicle: multiple two-tailed unpaired *t*-tests, total: *t*(8) = 4.6, *P* = 0.007; paw withdrawal: *t*(8) = 6.2, *P* = 0.001; paw shake: *t*(8) = 4.5, *P* = 0.007; paw lick and jump: *P* > 0.05; CP-55,940 vs. vehicle: multiple two-tailed unpaired *t*-tests, total: *t*(8) = 9.3, *P* < 0.0001; paw withdrawal: *t*(8) = 5.9, *P* = 0.002; paw shake, paw lick, and jump: *P* > 0.05, asterisks define differences to vehicle control after the Holm-Šídák multiple comparisons post-hoc test correction; **‘1350** vs. vehicle: two-way ANOVA; behavior x dose interaction: *F*(12, 80) = 8.2, *P* < 0.0001; behavior: *F*(4, 80) = 69.6, *P* < 0. 0001; dose: *F*(3, 80) = 34.2, *P* < 0.0001; asterisks define individual group differences to vehicle control after Dunnett’s multiple comparisons post-hoc test correction). **H.** Tail flick latency after co-treatment with the selective CB1 antagonist AM251 (all *n* = 5; one-way ANOVA, *F*(2, 17) = 29.9, *P* < 0.0001; asterisks define individual group differences to baseline control after Tukey’s multiple comparisons post-hoc test correction. **I.** Comparison of the effect of **‘4042** and CP-55,940 in wildtype (WT) versus CB2R knockout (KO) mice in the Hargreaves assay (all *n* = 5; two-way ANOVA; genotype x drug treatment interaction: *F*(2, 24) = 0.5, *P* > 0.05; genotype: *F*(1, 24) = 1.6, *P* > 0.05; drug treatment: *F*(2, 24) = 13.8, *P* = 0.0001; asterisks define individual group differences to baseline after Tukey’s multiple comparisons post-hoc test correction). **J.** Comparison of the effect of **‘4042** in wildtype (WT) versus CB2R knockout (KO) mice in the Tail Flick assay (all *n* = 5; two-way ANOVA; genotype x drug treatment interaction: *F*(1, 16) = 2.2, *P* > 0.05; genotype: *F*(1, 16) = 2.2, *P* > 0.05; drug treatment: *F*(1, 16) = 72.3, *P* < 0.0001; asterisks define individual group differences to baseline after Šídák’s multiple comparisons post-hoc test correction). **K.** Withdrawal latency in the Hargreaves assay after co-treatment with the selective CB2R antagonist SR 144528 (1 mg/kg) (all *n* = 5; one-way ANOVA, *F*(2, 17) = 6.6, *P* = 0.008; asterisks define individual group differences to vehicle control after Tukey’s multiple comparisons post-hoc test correction). **L.** Mesh grip test of catalepsy at 1 hour post-dose. Comparison of CP-55,940 (*n* = 5-10; one-way ANOVA, *F*(3, 26) = 10.3, *P* = 0.0001), haloperidol (*n* = 5; two-tailed unpaired *t*-test, *t*(8) = 3.5, *P* = 0.009), **‘4042** (*n* = 5; one-way ANOVA, *F*(3, 16) = 3.0, *P* > 0.05) and **‘1350** (*n* = 5-10; one-way ANOVA, *F*(3, 26) = 1.8, *P* > 0.05). Asterisks define differences between 1 mg/kg dose for each compound and respective vehicle control. Data at 30 min. timepoint are in Fig. 6. **M.** Comparison of morphine (*n* = 8; two-tailed unpaired t-test, *t*(14) = 2.51, *P* = 0.03) to CP-55,940 (*n* = 8; two-tailed unpaired t-test, *t*(14) = 2.9, *P* = 0.01) and **‘4042** (*n* = 8; two-tailed unpaired t-test, *t*(14) = 0.005, *P* > 0.05) in the Conditioned Place Preference (CPP) test. For all statistical tests: ns, not significant, \**P* < 0.05, \*\**P* < 0.01, \*\*\**P* < 0.001, \*\*\*\**P* < 0.0001. All data represent mean ± SEM of 5-10 animals.

## RESOURCE AVAILABILITY

### Materials availability

Compounds generated in this study can be purchased from Enamine.

### Data and code availability

The structure described in this manuscript were deposited to the Protein Data Bank under accession code 8GAG, and the map coordinates to EMDB under accession code EMD-29898. Additional data provided in the main text or supplemental materials. DOCK3.7 is freely available for non-commercial research in both executable and code form (http://dock.compbio.ucsf.edu/DOCK3.7/). A web-based version is freely available to all (http://blaster.docking.org/). The ultra-largelibrary used here is freely available (http://zinc15.docking.org, http://zinc20.docking.org). Any additional information required to reanalyze the data reported in this paper is available from the lead contact upon request.

## EXPERIMENTAL MODELS AND SUBJECT AVAILABILITY

### Cell lines

Sf9 cells were purchased from Expression Systems, Cat 94-001S.Tni Cells (Hi-5) were purchased from Expression Systems, Cat 94011S. HEK293 clonal cell line (HEK293SL cells) for bioSens-All experiments were derived from HEK293 cells purchased from ATCC. Rat brains were purchased from Bioivt, Cat. RAT00BRAINMZN. All cell lines are maintained by the supplier. No additional authentication was performed by the authors of this study. Sf9 cell lines were tested by the manufacturer for contamination, but not were not further tested by the authors of this study. HEK293 cells were tested for mycoplasma contamination on a regular basis. Cells were free of contaminations. Rat brains were not tested for mycoplasma contamination.

### Animal models

Behavioral testing was performed on adult (8-10 weeks old) C56BL/6 mice (strain #664 (male), strain #5786 (CB2R-deficient), or #36108 (CB1R - deficient) mice purchased from the Jackson Laboratory.

## METHODS

### Molecular docking

A crystal structure of the active-state CB1R receptor (PDB: 5XR8)^16^ was used for docking calculations. As the goal was to find small-molecule, non-phytocannabinoid ligands, we used ligand coordinates from the cryogenic ligand MDMB-Fubinaca (PDB: 6N4B)^18^, after overlaying the two receptor structures. The coordinates of Met363^6.55^ were modified slightly, while maintaining the residue within the electron density to reduce a clash with the overlaid ligand indole group. The combined coordinates were minimized with Schrӧdinger’s Maestro prior to calculation of the docking energy potential grids. These grids were precalculated using CHEMGRID^57^ for AMBER^58^ united atom van der Waals potential, QNIFFT^59^ for Poisson-Boltzmann-based electrostatic potentials, and SOLVMAP^60^ for Generalized Born-derived context-dependent ligand desolvation. Atoms of the ligand determined in the cryo-EM structure (PDB: 6N4B), MDMB-Fubinaca, were used to seed the matching sphere calculation in the orthosteric site, with 45 total spheres used (these spheres act as pseudo-atoms defining favorable sub-sites on to which library molecules may be superposed^61^. The receptor structure was protonated using REDUCE^62^ and AMBER united atom charges were assigned^58^. Control calculations^63^ using 324 known ligands extracted from the IUPHAR database^64^, CHEMBL24^33^, and ZINC15, and 14,929 property-matched decoys^65^ were used to optimize docking parameters based on enrichment measured by logAUC^63^, prioritization of neutral over charged molecules, and by the reproduction of expected and known binding modes of CB1R ligands. SPHGEN^61^ was used to generate pseudo-atoms to define the extended low protein dielectric and desolvation region^22,66^. The protein low dielectric and desolvation regions were extended as previously described^67^, based on control calculations, by a radius of 1.5 Å and 1.9 Å, respectively. The desolvation volume was removed around S383^7.39^ and H178^2.65^ to decrease the desolvation penalty near these residues and to increase the number of molecules that would form polar contacts with them.

A subset of 74 million large, relatively hydrophobic molecules from the ZINC15 database (http://zinc15.docking.org), with calculated octanol-water partition coefficients (cLogP, calculated using JChem-15.11.23.0, ChemAxon; https://www.chemaxon.com) between 3 and 5 and with molecular mass from 350 Da to 500 Da, was docked against the CB1R orthosteric site using DOCK3.7^68^. Of these, more than 18 million were successfully fit. An average of 4,706 orientations, and for each orientation, an average of 645 conformations was sampled. Overall, about 64 trillion complexes were sampled and scored. The total time was about 25,432 core hours, or less than 18 wall-clock hours on 1,500 cores.

To reduce redundancy of the top scoring docked molecules, the top 300,000 ranked molecules were clustered by ECFP4-based Tanimoto coefficient (Tc) of 0.5, and the best scoring member was chosen as the cluster representative molecule. These 60,420 clusters were filtered for novelty by calculating the Tc against >7,000 CB1R and CB2R receptor ligands from the CHEMBL24^33^ database. Molecules with Tc ≥ 0.38 to known CB1R/CB2R ligands were not pursued further.

After filtering for novelty, the docked poses of the best-scoring members of each cluster were filtered by the proximity of their polar moieties to Ser383^7.39^, Thr201^3.37^, or His178^2.65^, and visually inspected for favorable geometry and interactions. For the most favorable molecules, all members of its cluster were also inspected, and one of these was chosen to replace the cluster representative if they exhibited more favorable poses or chemical properties. Ultimately, 60 compounds were chosen for synthesis and testing.

### Make-on-demand synthesis and purity information

Of these 60, 52 were successfully synthesized by Enamine (an 87% fulfilment), but only 46 were ultimately screened due to poor DMSO solubility of six of the molecules. The purities of active molecules and analogs synthesized by Enamine were at least 90% and typically above 95%. The purity of compounds tested *in vivo* were >95% and typically above 98%. Synthetic routes^69^, chemical characterization, and purity quality control information for a subset of hits can be found in the supplementary information file and a list of all tested molecules and their single point displacement data can be found in **Supplementary Table 9**.

### Ligand optimization

Analogs with ECFP4 Tcs ≥ 0.5 to the four most potent docking hits (**‘51486**, **‘0450**, **‘7800**, and **‘7019**) were queried in Arthor and SmallWorld (https://sw.docking.org, https://arthor.docking.org; NextMove Software, Cambridge UK) against 1.4 and 12 Billion tangible libraries, respectively, the latter primarily containing Enamine REAL Space compounds (https://enamine.net/compound-collections/real-compounds/real-space-navigator). Results were pooled, docked into the CB1R site, and filtered using the same criteria as the original screen. Between 11 and 30 analogs were synthesized for each of the four scaffolds. Second- and third-round analogs were designed in 2D space based on specific hypotheses and were synthesized at Enamine.

### Radioligand Binding Experiments

The binding affinities of the compounds were obtained by competition binding using membrane preparations from rat brain (source of CB1) or HEK293 cells stably expressing human CB2R receptors and [^3^H]-CP-55,940 as the radioligand, as described^70^. The results were analyzed using nonlinear regression to determine the IC_50_ and K_i_ values for each ligand (Prism by GraphPad Software, Inc., San Diego, CA). The K_i_ values are expressed as the mean of two to three experiments each performed in triplicate.

### Functional assays

#### Lance Ultra cAMP Accumulation Assay

The inhibition of forskolin-stimulated cAMP accumulation assays was carried out using PerkinElmer’s Lance Ultra cAMP kit following the manufacturer’s protocol. In brief, CHO cells stably expressing human CB1R were harvested by incubation with Versene (ThermoFisher Scientific, Waltham, MA) for 10 min, washed once with Hank’s Balanced Salt Solution, and resuspended in stimulation buffer at ∼200 cells/μL density. The ligands at eight different concentrations (0.001-10,000 nM) in stimulation buffer (5 μL) containing forskolin (2 μM final concentration) were added to a 384-well plate followed by the cell suspension (5 μL; ∼1000 cells/well). The plate was incubated for 30 min at room temperature. Eu-cAMP tracer (5 μL) and Ulight-anti-cAMP (5 μL) working solutions were then added to each well, and the plate was incubated at room temperature for an additional 60 min. Results were measured on a Perkin-Elmer EnVision plate reader. The EC_50_ values were determined by nonlinear regression analysis using Prism software (GraphPad Software, Inc., San Diego, CA) and are expressed as the mean of three experiments, each performed in triplicate.

#### Cerep cAMP Inhibition Assay

Compounds **‘4042** and **‘3737** were run through the Cerep HTRF cAMP assay for functional activity as agonists (catalog number 1744; Cerep, Eurofins Discovery Services; France). The hCB1 CHO-K1 cells are suspended in HBSS buffer (Invitrogen) complemented with 20 mM HEPES (pH 7.4), then distributed in microplates at a density of 5.103 cells/well in the presence of either of the following: HBSS (basal control), the reference agonist at 30 nM (stimulated control) or the test compounds. Thereafter, the adenylyl cyclase activator forskolin is added at a final concentration of 25 μM. Following 30 min incubation at 37°C, the cells are lysed, and the fluorescence acceptor (D2-labeled cAMP) and fluorescence donor (anti-cAMP antibody labeled with europium cryptate) are added. After 60 min at room temperature, the fluorescence transfer is measured at λex=337 nm and λem=620 and 665 nm using a microplate reader (Envison, Perkin Elmer). The cAMP concentration is determined by dividing the signal measured at 665 nm by that measured at 620 nm (ratio). The results are expressed as a percent of the control response to 10 nM CP-55,940. Each measurement was done in triplicate.

#### Glosensor cAMP Accumulation Assay

The GloSensor cAMP accumulation assay was performed as secondary validation assays (dose-response setup) as described in detail on the NIMH PDSP website at https://pdsp.unc.edu/pdspweb/content/PDSP%20Protocols%20II%202013-03-28.pdf. The results were analysed using GraphPad Prism 9.1. Each experiment was performed in triplicate and functional IC_50_ values were determined from the mean of three independent experiments.

#### TRUPATH BRET2 G_oA_ recruitment for CB2R

CB2 receptor was co-expressed with. G_oA_ dissociation BRET2 assays were performed as previously described with minor modifications^71^. In brief, HEK293T cells were co-transfected overnight with human CB2 receptor, G_αo_A-Rluc, G_β3_, and G_γ9_-GFP2 constructs. After 18–24 hours, the transfected cells were seeded into poly-L-lysine-coated 384-well white clear-bottom cell culture plates at a density of 15,000–20,000 cells and incubated with DMEM containing 1% dialyzed FBS, 100 U mL−1 of penicillin and 100 µg ml−1 of streptomycin for another 24 hours. The next day, the medium was aspirated and washed once with 20 µL of assay buffer (1× HBSS, 20 mM HEPES, 0.1% BSA, pH 7.4). Then, 20 µL of drug buffer containing coelenterazine 400a (Nanolight Technology) at 5 µM final concentration was added to each well and incubated for 5 minutes, followed by the addition of 10 µL of 3X designated drug buffer for 5 minutes. Then, 10 µL of 4X final concentrations of ligands were added for 5 minutes. Finally, the plates were read in PHERAstar FSX (BMG Labtech) with a 410-nm (RLuc8-coelenterazine 400a) and a 515-nm (GFP2) emission filter, at 0.6-second integration times. BRET ratio was computed as the ratio of the GFP2 emission to RLuc8 emission. Data were normalized to percentage of CP-55,940 and analyzed in GraphPad Prism 9.1. Each experiment was performed in triplicate and functional IC_50_ values were determined from the mean of four independent experiments.

#### Tango β-arrestin-2 Recruitment Assay

The Tango β-arrestin-2 recruitment assays were performed as described^72^. In brief, HTLA cells were transiently transfected with human CB1R or CB2R Tango DNA construct overnight in DMEM supplemented with 10 % FBS, 100 µg ml−1 streptomycin and 100 U ml−1 penicillin. The transfected cells were then plated into poly-L-lysine-coated 384-well white clear-bottom cell culture plates in DMEM containing 1% dialysed FBS at a density of 10,000–15,000 cells per well. After incubation for 6 h, the plates were added with drug solutions prepared in DMEM containing 1% dialysed FBS for overnight incubation. On the day of assay, medium and drug solutions were removed and 20 µl per well of BrightGlo reagent (Promega) was added. The plates were further incubated for 20 min at room temperature and counted using the Wallac TriLux Microbeta counter (PerkinElmer). The results were analysed using GraphPad Prism 9.1. Each experiment was performed in triplicate and functional IC_50_ values were determined from the mean of three independent experiments.

#### DiscoverX PathHunter^®^ β-arrestin-2 Recruitment Assay

**‘4042** and **‘3737** were run through the PathHunter® β-arrestin-2 assay (catalog number 86-0001P-2070AG; DiscoverX, Eurofins Discovery Services; CA, USA). PathHunter cell lines (CHO-K1 lineage expressing hCB1) were expanded from freezer stocks according to standard procedures. Cells were seeded in a total volume of 20 μL into white walled, 384-well microplates and incubated at 37°C for the appropriate time prior to testing. For agonist determination, cells were incubated with sample to induce response. Intermediate dilution of sample stocks was performed to generate 5X sample in assay buffer. 5 μL of 5X sample was added to cells and incubated at 37°C or room temperature for 90 to 180 minutes. Vehicle concentration was 1%. Assay signal was generated through a single addition of 12.5 or 15 μL (50% v/v) of PathHunter Detection reagent cocktail, followed by a 1-hour incubation at room temperature. Microplates were read following signal generation with a PerkinElmer EnvisionTM instrument for chemiluminescent signal detection. Compound activity was analyzed using CBIS data analysis suite (ChemInnovation, CA). Percentage activity was calculated using the following equation:

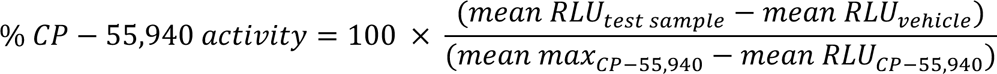

The data were analyzed in GraphPad Prism 9.1 using “dose–response-stimulation log(agonist) versus response (four parameters)” and data were presented as EC_50_ or pEC_50_ ± CIs of one independent experiment in duplicate.

#### Signaling profiling of hCB1 and hCB2 using bioSensAll^®^

ebBRET-based effector membrane translocation biosensor assays were conducted at Domain Therapeutics NA Inc. (Montreal, QC, Canada) as previously described^42^. CP-55,940, 2-AG and 25 test compounds were assayed for their effect on the signaling signature of the human cannabinoid receptor type 1 or 2 (hCB1 or hCB2) using the following bioSensAll^®^ sensors: the heterotrimeric G protein activation sensors (G_αs_, G_αi1_, G_αi2_, G_αoB_, G_αz_, G_α13_, G_αq_, G_α15_) and the ßarrestin-2 plasma membrane (PM) recruitment sensor (in the presence of GRK2 overexpression). HEK293 cells were maintained in Dulbecco’s Modified Eagle Medium (DMEM) (Wisent) supplemented with 1% penicillin-streptomycin (Wisent) and 10% (or 2 % for transfection) fetal bovine serum (Wisent) at 37°C with 5% CO2. All biosensor-coding plasmids and related information are the property of Domain Therapeutics NA Inc. The total amount of transfected DNA was adjusted and kept constant at 1 µg per mL of cell culture to be transfected using salmon sperm DNA (Invitrogen) as ‘carrier’ DNA, PEI (polyethylenimine 25 kDa linear, PolyScience) and DNA (3:1 ml PEI:mg DNA ratio) were first diluted separately in 150 mM NaCl then mixed and incubated for at least 20 minutes at room temperature to allow for the formation of DNA/PEI complexes. During the incubation, HEK293 cells were detached, counted, and re-suspended in maintenance medium to a 350,000 cells per mL density. At the end of the incubation period, the DNA/PEI mixture was added to the cells. Cells were finally distributed in 96-well plates (White Opaque 96-well /Microplates, Greiner) at a density of 35,000 cells per well. Forty-eight hours post-transfection, medium was aspirated and replaced with 100 µl of Hank’s Balanced Salt Solution buffer (HBSS) (Wisent) per well using 450-Select TS Biotek plate washer. After 60 min incubation in this medium, 10 µL of 10 µM e-Coelenterazine Prolume Purple (Methoxy e-CTZ) (Nanolight) was added to each well for a final concentration of 1 µM immediately followed by addition of increasing concentrations of the test compounds to each well using the HP D300 digital dispenser (Tecan). All compounds were assayed at 22 concentrations with each biosensor after a 10-minute room temperature incubation period. BRET readings were collected with a 0.4 sec integration time on a Synergy NEO plate reader (BioTek Instruments, Inc., USA; filters: 400nm/70nm, 515nm/20nm). BRET signals were determined by calculating the ratio of light emitted by GFP-acceptor (515nm) over light emitted by luciferase-donor (400nm). All BRET ratios were standardized using the universal BRET (uBRET) equation:

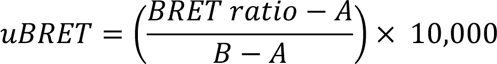

where *A* is the BRET ratio obtained from transfection of negative control and *B* is the BRET ratio obtained from transfection of positive control. Data were normalized to the best fit values of CP-55,940 from each individual experiment before being pooled across replicates. If CP-55,940 had no response, data were left unnormalized and *uBRET* was used for plotting. The data were analyzed using the four-parameter logistic non-linear regression model in GraphPad Prism 9.1 and data were presented as means ± CIs of 1-4 independent experiments.

For relative efficacy calculations for **‘1350** and **‘4042** versus CP-55940, first *E_max_* and *EC_50_* values were determined from dose-response curves to calculate the *log(E_max_/EC_50_)* value for each pathway and each compound. Then, the difference between the *log(E_max_/EC_50_)* values was calculated using the following equation:

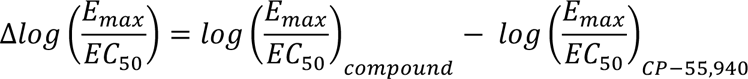

The SEM was calculated for the *log(E_max_/EC_50_)* ratios using the following equation:

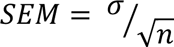

where σ is the standard deviation, and n is the number of experiments.

The SEM was calculated for the Δ*log(E_max_/EC_50_)* ratios using the following equation:

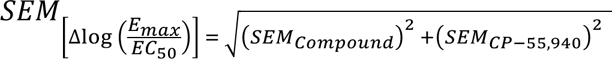

The compounds’ efficacy toward each pathway, relative to CP-55,940, were finally calculated using the following equation:

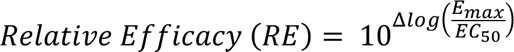

The relative efficacies were used in radar plots to demonstrate the relative compound effectiveness compared to CP-55,940.

Statistical analysis was performed using a two-tailed unpaired t-test on the Δ*log(E_max_/EC_50_)* ratios to make pairwise comparisons between tested compounds and CP-55,940 for a given pathway, where P < 0.05 was considered statistically significant.

#### Bimane Fluoroscence

A minimal cysteine version of CB1R was generated^73^ where all the cysteine residues (except C256 and C264) were mutated to alanine. A cysteine residue was engineered at residue 336 (L6.28) on TM6, which was labeled with monobromobimane (bimane) by incubating 10 μM receptor with 10-molar excess of bimane at room temperature for one hour. Excess label was removed using size exclusion chromatography on a Superdex 200 10/300 Increase column in 20 mM HEPES pH 7.5, 100 mM NaCl and 0.01% MNG/0.001% CHS. Bimane-labeled CB1R at 0.1 mM was incubated with ligands (10 μM) for one hour at room temperature. Fluorescence data was collected at room temperature in a 150 μL cuvette with a *FluorEssence v3.8 software* on a Fluorolog instrument (*Horiba*) in photon-counting mode. Bimane fluorescence was measured by excitation at 370 nm with excitation and emission bandwidth passes of 4 nm. The emission spectra were recorded from 410 to 510 nm with 1 nm increment and 0.1 s integration time.

#### GTP turnover assay

Analysis of GTP turnover was performed by using a modified protocol of the GTPase-Glo^TM^ assay (Promega) described previously^74^. Ligand-bound (10 μM ligand incubated for one hour at room temperature) or apo CB1R (1 μM) was mixed with G-protein (1 μM) in 20 mM HEPES, pH 7.5, 50 mM NaCl, 0.01% L-MNG/0.001% CHS, 100 μM TCEP, 10 μM GDP and 10 μM GTP and incubated at room temperature. GTPase-Glo-reagent was added to the sample after incubation for 60 minutes (G_i1-3_) and 20 minutes for (G_o_). Luminescence was measured after the addition of detection reagent and incubation for 10 min at room temperature using a *SpectraMax Paradigm* plate reader.

### Colloidal Aggregation Counter-Screens

#### Dynamic Light Scattering (DLS)

Samples were prepared as 8-point half-log dilutions in filtered 50 mM KPi buffer, pH 7.0 with final DMSO concentration at 1% (v/v). Colloidal particle formation was measured using DynaPro Plate Reader II (Wyatt Technologies). All compounds were screened in triplicate.

#### Enzyme Inhibition Counter-Screening Assays

Enzyme inhibition assays to test for colloidal inhibition were performed at room temperature using CLARIOstar Plate Reader (BMG Labtech). Samples were prepared in 50 mM KPi buffer, pH 7.0 with final DMSO concentration at 1% (v/v). Compounds were incubated with 2 nM AmpC β-lactamase (AmpC) or Malate dehydrogenase (MDH) for 5 minutes. AmpC reactions were initiated by the addition of 50 μM CENTA chromogenic substrate (219475, Calbiochem). The change in absorbance was monitored at 405 nm for CENTA (219475, Calbiochem) or 490 for Nitrocefin (484400, Sigma Aldrich) for 60 sec. MDH reactions were initiated by the addition of 200 μM nicotinamide adenine dinucleotide (NADH) (54839, Sigma Aldrich) and 200 μM oxaloacetic acid (324427, Sigma Aldrich). The change in absorbance was monitored at 340 nm for 60 sec. Initial rates were divided by the DMSO control rate to determine % enzyme activity. Each compound was screened at 100μM in triplicate for three independent experiments, if enzyme inhibition greater than 30% was observed, 8-point half-log concentrations were performed in triplicate for three independent experiments. Data was analyzed using GraphPad Prism software version 9.1 (San Diego, CA).

### Cryo-EM sample preparation and structure determination

#### Purification of hCB1

hCB1R was expressed and purified as described previously^18^. An N-terminal FLAG tag and C-terminal histidine tag was added to human full-length CB1. This CB1R construct was expressed in *Spodoptera frugiperda Sf9* insect cells with the baculovirus method (Expression Systems). Insect cell pellets expressing CB1R was solubilized with buffer containing 1% lauryl maltose neopentyl glycol (L-MNG) and 0.1% cholesterol hemisuccinate (CHS) and purified by nickel-chelating Sepharose chromatography. The Ni column eluant was applied to a M1 anti-FLAG immunoaffinity resin. After washing to progressively decreasing concentration of L-MNG, the receptor was eluted in a buffer consisting of 20 mM HEPES pH 7.5, 150 mM NaCl, 0.05% L-MNG, 0.005% CHS, FLAG peptide and 5 mM EDTA. As the final purification step, CB1R was applied to a Superdex 200 10/300 gel filtration column (GE) in 20 mM HEPES pH 7.5, 150 mM NaCl, 0.02% L-MNG, 0.002% CHS. Ligand-free CB1R was concentrated to ∼500 µM and stored in −80 °C.

#### Expression and purification of G_i/o_ heterotrimer

Expression and purification of all heterotrimeric G-protein (G_i/o_) follow similar protocols. Heterotrimeric G_i_ was expressed and purified as previously described^75^. Wild-type human Gα_i1_ subunit virus and wild-type human β_1_ψ_2_ (with histidine tagged β subunit) virus were used to co-infect Insect (*Trichuplusia ni, Hi5*) cells. Cells expressing the heterotrimetric, G_i_β_1_ψ_2_ G-protein were lysed in hypotonic buffer and G-protein was extracted in a buffer containing 1% sodium cholate and 0.05% n-dodecyl-β-D-maltoside (DDM, Anatrace). Detergent was exchanged from cholate/DDM to DDM on Ni Sepharose column. The eluant from the Ni column was dialyzed overnight into 20 mM HEPES, pH 7.5, 100 mM sodium chloride, 0.1% DDM, 1 mM magnesium chloride, 100 μM TCEP and 10 μM GDP together with Human rhinovirus 3C protease (3C protease) to cleave off the His tag in the β subunit. 3C protease was removed by Ni-chelating sepharose and the heterotrimetric G-protein was further purified with MonoQ 10/100 GL column (GE Healthcare). Protein was bound to the column and washed in buffer A (20 mM HEPES, pH 7.5, 50 mM sodium chloride, 1 mM magnesium chloride, 0.05% DDM, 100 μM TCEP, and 10 μM GDP). The protein was eluted with a linear gradient of 0–50% buffer B (buffer A with 1 M NaCl). The collected G protein was dialyzed into 20 mM HEPES, pH 7.5, 100 mM sodium chloride, 1 mM magnesium chloride, 0.02% DDM, 100 μM TCEP, and 10 μM GDP. Protein was concentrated to about 200 µM and flash frozen until further use.

#### Purification of scFv16

scFv16 was purified with a hexahistidine-tag in the secreted form from *Trichuplusia ni Hi5* insect cells using the baculoviral method. The supernatant from baculoviral infected cells was pH balanced and quenched with chelating agents and loaded onto Ni resin. After washing with 20 mM HEPES pH 7.5, 500 mM NaCl, and 20 mM imidazole, protein was eluted with 250 mM imidazole. Following dialysis with 3C protease into a buffer consisting of 20 mM HEPES pH 7.5 and 100 mM NaCl, scFv16 was further purified by reloading over Ni a column. The collected flow-through was applied onto a Superdex 200 16/60 column and the peak fraction was collected, concentrated and flash frozen.

#### CB1-G_i1_ complex formation and purification

CB1R in L-MNG was incubated with excess **‘1350** for ∼1 hour at room temperature. Simultaneously, G_i1_ heterotrimer in DDM was incubated with 1% L-MNG/0.1% CHS at 4 °C. The **‘1350**-bound CB1R was incubated with a 1.25 molar excess of detergent exchanged G_i_ heterotrimer at room temperature for ∼ 3 hour. The complex sample was further incubated with apyrase for 1.5 hour at 4 °C to stabilize a nucleotide-free complex. 2 mM CaCl_2_ was added to the sample and purified by M1 anti-FLAG affinity chromatography. After washing to remove excess G protein and reduce detergents, the complex was eluted in 20mM HEPES pH 7.5, 100mM NaCl, 0.01% L-MNG/0.001% CHS, 0.0033% GDN/0.00033% CHS, 10 µM **‘1350**, 5 mM EDTA, and FLAG peptide. The complex was supplemented with 100 µM TCEP and incubated with 2 molar excess of scFv16 overnight at 4 °C. Size exclusion chromatography (Superdex 200 10/300 Increase) was used to further purify the CB1-G_i_-scFv16 complex. The complex in 20mM HEPES pH 7.5, 100mM NaCl, 10 µM **‘1350**, 0.00075% L-MNG/0.000075% CHS and 0.00025% GDN/0.000025% CHS was concentrated to ∼12 mg/mL for electron microscopy studies.

#### Cryo-EM data acquisition

Grids were prepared by applying 3 μL of purified CB1-G_i_ complex at 12 mg/ml to glow-discharged holey carbon gold grids (Quantifoil R1.2/1.3, 200 mesh). The grids were blotted using a Vitrobot Mark IV (FEI) with 3 s blotting time and blot force 3 at 100% humidity at room temperature and plunge-frozen in liquid ethane. A total of 8324 movies were recorded on a Titan Krios electron microscope (Thermo Fisher Scientific-FEI) operating at 300 kV at a calibrated magnification of 96,000x corresponding to a pixel size of 0.8521 Å. Micrographs were recorded using a K3 Summit direct electron camera (Gatan Inc.) with a dose rate of 16.4 electrons/pixel/s. The total exposure time was 2.5 s with an accumulated dose of ∼ 56.6 electrons per Å^2^ and a total of 50 frames per micrograph. Automatic data acquisition was done using *SerialEM*.

#### Image processing and 3D reconstructions

Micrographs were subjected to beam-induced motion correction using *MotionCor2*^76^ implemented in Relion 2.1.0^77^. CTF parameters for each micrograph were determined by *CTFFIND4*^78^. An initial set of 4,967,593 particle projections were extracted using semi-automated procedures and subjected to reference-free two-dimensional and multiple rounds of three-dimensional classification in *Relion 2.1.0*^77^ to remove low-resolution and otherwise poor-quality particles. From this step, 750,496 particle projections were selected for further processing in *CryoSPARC*^79^. A final two-dimensional classification step in order to select for the highest-resolution particles resulted in a particle set containing 465,411 particles. These particles were reconstructed to a global nominal resolution of 3.3 Å (**Fig. S5**) at FSC of 0.143 using non-uniform refinement. Local resolution was estimated within *CryoSPARC*^79^.

#### Model building and refinement

The initial template of CB1R was the MDMB-Fubinaca-bound CB1-G_i_ complex structure (PDB: 6N4B). *Phenix.elbow was used to generate* Agonist coordinates and geometry restrains. Models were docked into the EM density map using *UCSF Chimera*. *Coot* was used for iterative model building and the final model was subjected to global refinement and minimization in real space using *phenix.real_space_refine* in *Phenix*. Model geometry was evaluated using *Molprobity*. FSC curves were calculated between the resulting model and the half map used for refinement as well as between the resulting model and the other half map for cross-validation (**Fig. S5**). The final refinement parameters are provided in **Supplementary Table 3**. The ligand symmetry accounted RMSD between the docked pose and cryo-EM pose of **‘1350** was calculated by the Hungarian algorithm in DOCK6^80^.

### Off-target activity

#### GPCRome and Comprehensive Binding Panel

Compound **‘4042** was tested at 10 μM for off-target activity against a panel of 320 non-olfactory GPCRs using PRESTO-Tango GPCRome arrestin-recruitment assay, as described^72^. Receptors with at least three-fold increased relative luminescence over corresponding basal activity are potential positive hits, and were tested in dose response follow-up studies. Compound **‘4042** was further tested at 1 µM for off-target activity at a panel of 45 common GPCR and non-GPCR drug targets. Receptors with at least 50% displaced radioligand are potential positive hits and were tested in dose response follow-up studies. Screening was performed by the National Institutes of Mental Health Psychoactive Drug Screen Program (PDSP)^81^. Detailed experimental protocols are available on the NIMH PDSP website at https://pdsp.unc.edu/pdspweb/content/PDSP%20Protocols%20II%202013-03-28.pdf.

### In vivo methods

#### Animals and ethical compliance

Animal experiments were approved by the UCSF Institutional Animal Care and Use Committee and were conducted in accordance with the NIH Guide for the Care and Use of Laboratory animals (protocol #AN195657). Adult (8-10 weeks old) male C56BL/6 (strain # 664), CB1R knockout (strain #36108), and CB2R knockout (strain #5786) mice were purchased from the Jackson Laboratory. Mice were housed in cages on a standard 12:12 hour light/dark cycle with food and water ad libitum. Sample sizes were modelled on our previous studies and on studies using a similar approach, which were able to detect significant changes^82,83^. The animals were randomly assigned to treatment and control groups. Animals were initially placed into one cage and allowed to freely run for a few minutes. Then each animal was randomly picked up, injected with compound treatment or vehicle, and placed into a separate cylinder before the behavioral test.

#### *In vivo* compound preparation

Ligands were sourced from Enamine (**‘4042**) or Sigma-Aldrich (CP-55,940, Cat No. C1112; Haloperidol, Cat. No. H1512; AM251, Cat. No. A6226; SR 144528, Cat. No. SML1899) and dissolved 30 min before injections. **‘4042** was resuspended in a 20% Kolliphor HS-15 (Sigma-Aldrich, Cat. No. 42966) / 40% saline / 40% water for injections (v/v/v) vehicle for i.p. injections. CP-55,940, SR 144528, and AM251 for i.p. injections and **‘4042** for i.t. injections were resuspended in a 5% EtOH /5% Kolliphor-EL (Sigma-Aldrich Cat. No. C5135) / 90% water for injections vehicle. Morphine (provided by the NIH) was resuspended in 100% saline. Haloperidol was resuspended in 20% cyclodextrin (Sigma-Aldrich, Cat. No. H107). All cannabinoid formulations were prepared in silanized glass vials.

#### Pharmacokinetics

Pharmacokinetic experiments were performed by Bienta (Enamine Biology Services) in accordance with Enamine pharmacokinetic study protocols and Institutional Animal Care and Use Guidelines (protocol number 1-2/2020). Plasma, brain, and CSF concentrations were measured for **‘4042** and CP-55,940 following a 0.2 mg/kg intraperitoneal (i.p.) dose. The batches of working formulations were prepared 5-10 minutes prior to the *in vivo* study. In each compound study, up to nine time points (5, 15, 30, 60, 120, 240, 360, 480 and 1440 min) were collected; each of the time point treatment groups included 3 male CD-1 mice. There was also a one mouse control group. All animals were fasted for 4 h before dosing. Mice were injected i.p. with 2,2,2-tribromoethanol at the dose of 150 mg/kg prior to drawing CSF and blood. Blood collection was performed from the orbital sinus in microtainers containing K_3_EDTA. CSF was collected under a stereomicroscope from cisterna magna using 1 mL syringes. Animals were sacrificed by cervical dislocation after the blood samples collection. After this, right lobe brain samples were collected and weighted. All samples were immediately processed, flash-frozen and stored at −70°C until subsequent analysis.

Plasma samples (40 μL) were mixed with 200 μL of internal standard solution. After mixing by pipetting and centrifuging for 4 min at 6,000 rpm, supernatant was injected into LC-MS/MS system. Solution of Difenoconazole (50 ng/ml in water-methanol mixture 1:9, v/v) was used as the internal standard (IS) for quantification of **‘4042** and mefenamic acid (100 ng/mL in water-acetonitrile mixture 1:9, v/v) was used as the IS for the quantification of CP-55,940. Brain samples (weight 59 mg – 201 mg) were homogenized with 5 volumes of IS(80) solution using zirconium oxide beads (115 mg ± 5 mg) in The Bullet Blender® homogenizer for 30 seconds at speed 8. After this, the samples were centrifuged for 4 min at 14,000 rpm, and supernatant was injected into LC-MS/MS system. CSF samples (4 μL) were mixed with 100 μL of IS(80) solution. After mixing by pipetting and centrifuging for 4 min at 6,000 rpm, 1-6 μL of each supernatant was injected into LC-MS/MS system.

Analyses of plasma, brain and CSF samples were conducted at Enamine/Bienta. The concentrations of compounds in samples were determined using high performance liquid chromatography/tandem mass spectrometry (HPLC-MS/MS) method. Data acquisition and system control was performed using Analyst 1.6.3 software (AB Sciex, Canada). The concentrations of the test compound below the lower limit of quantitation (LLOQ: 2-5 ng/mL for plasma and CSF, 1-5 ng/g for brain) were designated as zero. The pharmacokinetic data analysis was performed using noncompartmental, bolus injection or extravascular input analysis models in WinNonlin 5.2 (PharSight). Data below LLOQ were presented as missing to improve validity of T½ calculations.

#### Behavioral analyses

For all behavioral tests, the experimenter was always blind to treatment. Animals were first habituated for 30-60 minutes in Plexiglas cylinders and then tested 30 minutes after i.p. or i.t. injection of the compounds. The mechanical (von Frey), thermal (Hargreaves, hotplate and tail flick) and ambulatory (rotarod) tests were conducted as described^84^. Hindpaw mechanical thresholds were determined with von Frey filaments using the up-down method^85^. Hindpaw thermal sensitivity was measured with a radiant heat source (Hargreaves) or on a hotplate at 52°C. For the tail flick assay, sensitivity was measured by immersing the tail into a 50°C water bath. For the ambulatory (rotarod) test, before testing with any compound, mice underwent three trainings on three consecutive days (until they reach 300 sec). Each training has three sessions of five min. each. Therapeutic index was calculated as the ratio of the minimum dose of side effect phenotype and the minimum dose of analgesic phenotype.

#### SNI model of neuropathic pain

Under isoflurane anesthesia, two of the three branches of the sciatic nerve were ligated and transected distally^86^, leaving the sural nerve intact. Behavior was tested 7 to 14 days after injury.

#### CFA

The CFA model of chronic inflammation was induced as described previously^87^. Briefly, CFA (Sigma) was diluted 1:1 with saline and vortexed for 30 min. When fully suspended, we injected 20 μL of CFA into one hindpaw. Heat thresholds were measured before the injection (baseline) and 3 days after the injection using the Hargreaves test.

#### Open Field Test

Thirty minutes after i.p. injection, mice were placed in the center of a round open-field (2 feet diameter) and their exploratory behavior recorded over the next 15 minutes. Distance traveled was used to represent open field behavior.

#### Conditioned Place Preference

To determine if **‘4042** was inherently rewarding or aversive we used the conditioned place paradigm as described^88^. Briefly, mice were first habituated to the test apparatus, twice, and their preference for each chamber recorded for 30 minutes (Pretest). Two conditioning days followed in which mice received the vehicle control or the compound, and 30 minutes later restricted for 30 minutes in the preferred or non-preferred chamber, respectively. On day 5 (Test day), mice were allowed to roam freely between the 3 chambers of the apparatus and their preference for each chamber recorded for 30 minutes. To calculate the CPP score, we subtracted the time spent in each chamber of the box on the Pretest day from that of the Test day (CPP score = Test - Pretest).

#### Acetone Test

Mice were placed on a wire mesh and thirty min after an i.p. injection of the compounds we applied a drop (50 μL) of acetone on the ventral aspect of the hindpaw, 5 times every 30 sec. We recorded the number of nocifensive behaviors (paw lifts/licks/shakes/bites) over the 5 applications.

#### Formalin Test

Thirty minutes after an i.p. injection of the compounds, mice received an intraplantar injection of a 20μl solution containing 2% formalin (Acros Organics) and we recorded the time mice spent licking/biting/guarding (nocifensive behaviors) the injected hindpaw over the next 60 min.

#### Catalepsy Test

Thirty and 60 minutes after an i.p. injection of the compounds, mice were placed on a vertical wire mesh and the latency to move all four paws was recorded.

#### Body temperature measurements

Body temperature (BT) was measured using a telemetric probe device (HD-X10; Data Science International). Briefly, under anesthesia, the probe device was placed in the mouse abdomen and a subcutaneous tunnel was created from the neck to the abdominal skin, through which a catheter (connected to the probe) was pulled and then inserted into the left carotid artery. Three weeks later, the implanted mice were singly housed in a cage that was placed on top of the DSI receiver (for probe signal detection). We monitored BT continuously over 2h, in the following manner: 30 minutes (for baseline), 30 minutes after injection of the vehicle and then for 1h after injection of the compound. Data were acquired using the Ponemah Telemetry acquisition software (DSI) and percent changes were presented relative to each mouse’s baseline.

## STATISTICAL ANALYSIS

### Statistical analyses

All statistical tests were run with GraphPad Prism 9.1 (GraphPad Software Inc., San Diego). A two-tailed unpaired *t*-test was used to compare the pKi ± 95% CI for **‘4042** at CB1R versus CB2R (**Fig. S8** legend). Experiments of the compounds in the in vivo assays were analyzed by unpaired two-tailed t-tests, one-way ANOVA, or two-way ANOVA, depending on the experimental design. All statistical calculations were controlled for multiple hypothesis testing using a post-hoc test as described in the **Fig. 5**, **Fig. 6**, or **Fig. S11** legends. Details of the analyses, including groups compared in post-hoc sets, number of animals per group, *t* or *F* statistics, *P* values, definition of center, and dispersion and precision measures can be found in the figure legends. The animals were randomly assigned to the treatment group and control group. For behavioral experiments, animals were initially placed into one cage and allowed to free run for a few minutes. Next, each animal was randomly picked up, injected with the drug or vehicle control and placed into a separate cylinder before the behavioral test. Explicit sample size calculations were not performed but were instead modeled on previous studies using a similar approach which was demonstrated to be capable of detecting significant changes.

