## Supplementary Information for "Large library docking for cannabinoid-1 receptor agonists with reduced side effects"

This file includes:

Supplementary Tables 1 to 8

Additional supplementary materials:

Synthetic procedures, chemical characterization and spectral data is supplied as
Supplemental_Information_QC.pdf
Supplementary_Table_9.xlsx
Model_final.pdb
Map_final.mrc
PDB_Validation_report.pdf

**Supplementary Table 1. Binding affinities for hits identified in initial CB1 docking screen.**

| Compound | Global rank | rCB1 affinity[a]<br>$K_i$ [95% CI] nM<br>$pK_i$ [95% CI] | Tc[b] | Nearest ChEMBL ligand[c] |
|---|---|---|---|---|
| 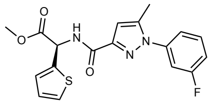<br>‘51486 | 117390 | 731 [552 – 969]<br>6.14 [6.01 –6.26] | 0.30 | 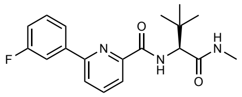<br>CHEMBL4110127 |
| 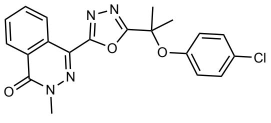<br>‘0450 | 6582 | 691 [459 – 1033]<br>6.16 [5.99 – 6.34] | 0.36 | 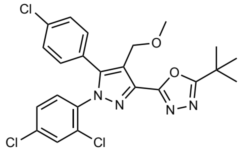<br>CHEMBL519214 |
| 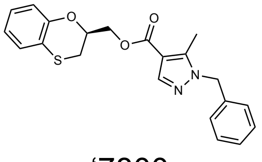<br>‘7800 | 12210 | 1007 [615 – 1654]<br>6.0 [5.78 – 6.21] | 0.28 | 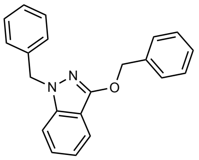<br>CHEMBL3116279 |
| 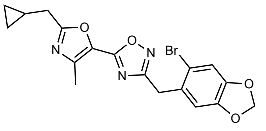<br>‘7019 | 20488 | 4039 [3027 – 5379]<br>5.39 [5.27 –5.52] | 0.24 | 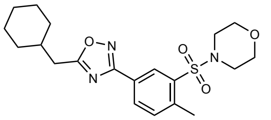<br>CHEMBL472680 |
| 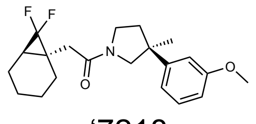<br>‘7218 | 29322 | 52.2% [24.79] | 0.31 | 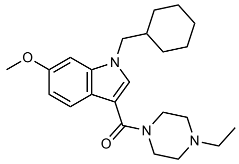<br>CHEMBL3347301 |
| 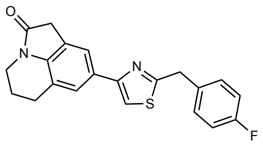<br>‘1038 | 47606 | 53.6% [2.91] | 0.28 | 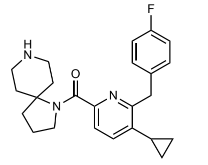<br>CHEMBL3890211 |
| 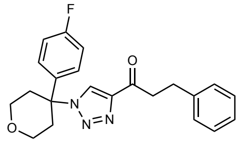<br>‘7337 | 24720 | 57.0% [3.04] | 0.29 | 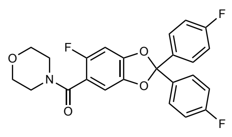<br>CHEMBL259699 |
| 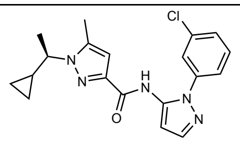<br>‘7902 | 139929 | 57.1% [0.02] | 0.31 | 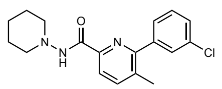<br>CHEMBL3915046 |

| | | | | |
|---|---|---|---|---|
| 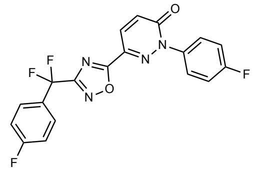 '2443 | 21964 | 51.1% [4.87] | 0.23 | 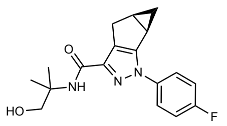 CHEMBL3354970 |

[a]Binding affinity to rCB1 represented as Ki [95% CI] and pKi [95% CI] from three independent experiments in triplicate when measured. Otherwise, % radioligand displacement [S.D] from three replicates in a single-point competition experiment at 10 μM

[b]Tanimoto coefficient based on ECFP4 fingerprints

[c]Corresponding ChEMBL ligand with the most similar fingerprint

**Supplementary Table 2. Binding affinities and functional activities for active analogs at CB1.**

| Compound | rCB1 binding<br>$K_i$ [95% CI] (nM)<br>$pK_i$ [95% CI]<br>$E_{max}$ [SEM]<br>Significance[a] | hCB1 Lance Ultra cAMP<br>$EC_{50}$ [95% CI] (nM)<br>$pEC_{50}$ [95% CI]<br>$E_{max}$ [95% CI]<br>Significance[a] | hCB1 Cerep cAMP<br>$EC_{50}$ [95% CI] (nM)<br>$pEC_{50}$ [95% CI]<br>$E_{max}$ [95% CI] | hCB1 Glosensor cAMP<br>$EC_{50}$ [95% CI] (nM)<br>$pEC_{50}$ [95% CI]<br>$E_{max}$ [95% CI] | hCB1 Tango ß-arrestin recruitment<br>$EC_{50}$ [95% CI] (nM)<br>$pEC_{50}$ [95% CI]<br>$E_{max}$ [95% CI] | hCB1 DiscoverX ß-arrestin recruitment<br>$EC_{50}$ [95% CI] (nM)<br>$pEC_{50}$ [95% CI]<br>$E_{max}$ [95% CI] |
|---|---|---|---|---|---|---|
| 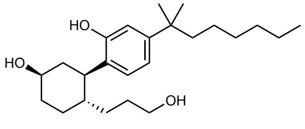<br>**CP–55,940** | 2.9 [2.05 – 4.2]<br>8.5 [8.4 – 8.7]<br>98% [3.5] | 6.2 [4.7 – 8.0]<br>8.2 [8.1 – 8.3]<br>85% [86 – 85] | 0.026 | 0.028 [0.02 – 0.04]<br>10.6 [10.5 – 10.7]<br>96% [93 – 99] | 8.9 [7.5 – 10.6]<br>8.1 [8.0 – 8.1]<br>100% [96 – 104] | 4.0 [3.2 – 4.9]<br>8.4 [8.3 – 8.5]<br>108% [99 – 109] |
| 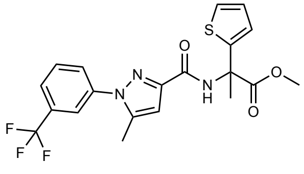<br>**‘4042** | 1.86 [1.37 – 2.52]<br>8.7 [8.6 – 8.9]<br>99% [3.0]<br>ns | 3.3 [1.9 – 5.6]<br>8.5 [8.3 – 8.7]<br>78% [78 – 79]<br>ns | 0.008 [0.006 – 0.01]<br>11.1 [11.0 – 11.2]<br>96% [102 – 107] | 0.039 [2.9 – 5.4]<br>10.4 [10.3 – 10.5]<br>91% [87 – 94] | 10.7 [8.7 – 13.3]<br>8.0 [7.9 – 8.1]<br>102% [98 – 105] | 2.3 [2.5 –4.8]<br>8.7 [8.3 – 9.6]<br>71% [60 – 65] |
| 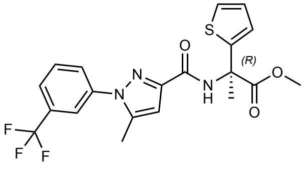<br>**‘1350** | 0.95[.74 – 1.24]<br>9.02 [8.9 – 9.1]<br>106% [2.9]<br>* | 1.6 [0.7 – 3.6]<br>8.8 [8.4 – 9.2]<br>78% [77 – 80]<br>** | -- | -- | -- | -- |
| 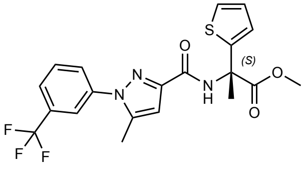<br>**‘8690** | 90.2 [56.7 – 143]<br>7.1 [6.9 – 7.3]<br>100% [4.0]<br>**** | 473 [109 – 1822]<br>6.3 [5.8 – 7.0]<br>53% [45 – 65]<br>**** | -- | -- | -- | -- |

| | | | | | | |
|---|---|---|---|---|---|---|
| 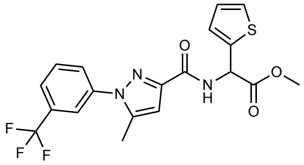 **‘60154** | 44.3 [33.9 – 58.0]<br>7.4 [7.2 – 7.5]<br>110% [2.9] | 351 [93]<br>6.5 [7.0]<br>67% [59 – 89] | -- | 25.2 [16 – 40]<br>7.6 [7.4 – 7.8]<br>82% [74 – 189] | 819 [718 – 934]<br>6.1 [6.0 – 6.1]<br>39% [37 – 40] | -- |
| 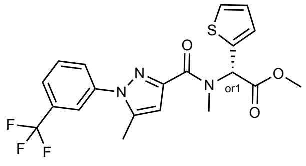 **‘1066** | 1719 [736 – 4048]<br>5.8 [5.4 – 6.1]<br>97% [8.0] | | | | | |
| 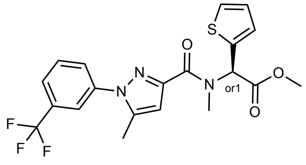 **‘6000** | 1455 [943 – 2249]<br>5.8 [5.7 – 6.0]<br>96% [4.0] | | | | | |
| 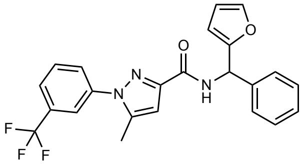 **‘1081** | 116 [76.3 – 178]<br>6.9 [6.8 – 7.1]<br>96% [4.0] | -- | -- | -- | -- | -- |
| 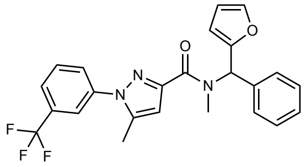 **‘1082** | 850 [488 – 1491]<br>6.1 [5.8 – 6.3]<br>94% [5.7] | -- | -- | -- | -- | -- |

| | | | | | | |
|---|---|---|---|---|---|---|
| 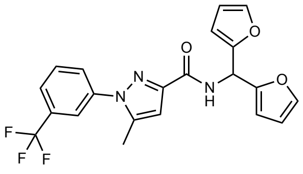<br>**‘1090** | 90.8 [42.7 – 192]<br>7.0 [6.7 – 7.4]<br>99% [7.13] | -- | -- | -- | -- | -- |
| 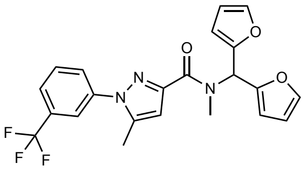<br>**‘4388** | 1360 [998 – 1857]<br>5.9 [5.7 – 6.0]<br>113% [4.4] | -- | -- | -- | -- | -- |
| 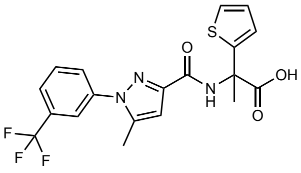<br>**‘4051** | 5328 [3774– 7507]<br>5.3 [5.1 – 5.4]<br>103% [3.8] | -- | -- | -- | -- | -- |
| 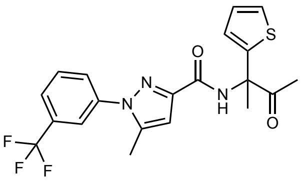<br>**‘4156** | 13.5 [6.3 – 30.1]<br>7.9 [7.5 – 8.2]<br>91% [7.5] | | | | | |
| 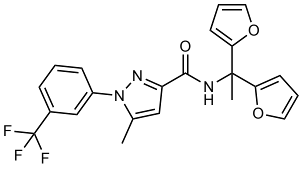<br>**‘4936** | 7.5 [3.9 – 14.3]<br>8.1 [7.9 – 8.4]<br>88% [6.3] | | | | | |

| | | | | | | |
|---|---|---|---|---|---|---|
| 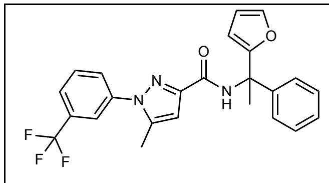<br>**‘5806** | 8.0 [3.0 – 21.8]<br>8.1 [7.7 – 8.5]<br>100% [10.5] | | | | | |
| 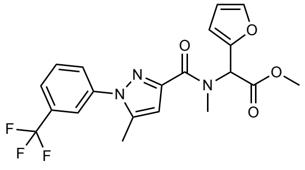<br>**‘6425** | 934 [583 – 1501]<br>6.0 [5.8 – 6.2]<br>108% [5.8] | -- | -- | -- | -- | -- |
| 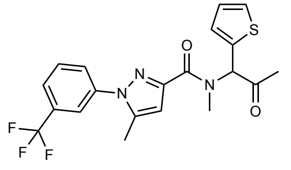<br>**‘6829** | 1046 [669 –<br>1643]<br>6.0 [5.8 – 6.2]<br>106% [5.8] | -- | -- | -- | -- | -- |
| 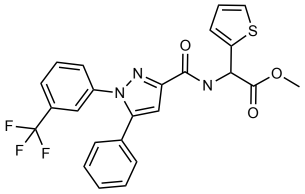<br>**‘8079** | 18.5 [13.8 – 25.0]<br>7.7 [7.6 – 7.9]<br>104% [3.0] | -- | -- | -- | -- | -- |
| 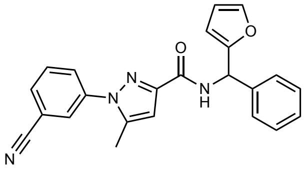<br>**‘12565** | 301 [195 – 462]<br>6.5 [6.3 – 6.7]<br>97% [4.0] | -- | -- | -- | -- | -- |

| | | | | | | |
|---|---|---|---|---|---|---|
| 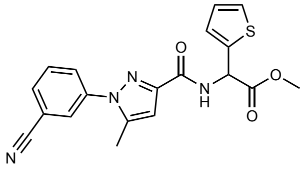 **‘31604** | 801 [596 –1,076]<br>6.1 [6.0 – 6.2]<br>100% [2.8] | -- | -- | -- | -- | -- |
| 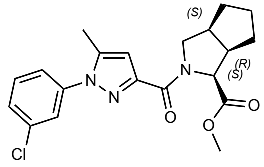 **‘10010** | 1,196 [952 – 1,505]<br>5.9 [5.8 – 6.0]<br>101% [2.9] | -- | -- | N.D | > 10,000 | -- |
| 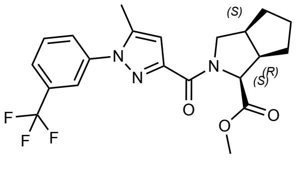 **‘6439** | 251 [173 – 364]<br>6.6 [6.4 – 6.8]<br>106% [3.9] | -- | -- | N.D. | > 10,000 | -- |
| 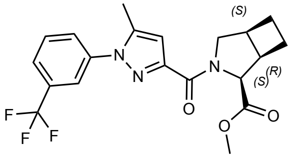 **‘6448** | 866 [564 – 1,317]<br>6.1 [5.9 – 6.3]<br>103% [4.1] | -- | -- | -- | -- | -- |
| 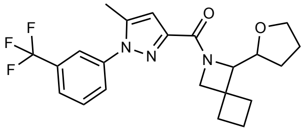 **‘3737** | 173 [94 – 322]<br>6.76 [6.5 – 7.03]<br>112% [8.8] | -- | 326 [168 – 1044]<br>6.5 [6.0 – 6.8]<br>99% [91 – 135] | N.D | > 10,000 | > 10,000 |
| 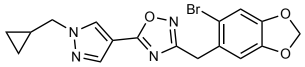 **‘5424** | 876 [683 – 1123]<br>6.1 [6.0 – 6.2]<br>104% [2.6] | -- | -- | -- | -- | -- |

| | | | | | | |
|---|---|---|---|---|---|---|
| 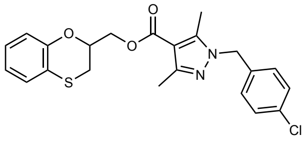 **'5463** | 825 [396 – 1,755]<br>6.1 [5.8 – 6.4]<br>88% [6.6] | -- | -- | -- | -- | -- |
| 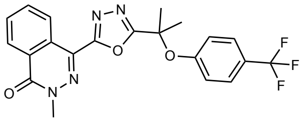 **'2153** | 163 [90 – 287]<br>6.79 [6.5 – 7.0]<br>82.6% [4.5] | -- | -- | -- | -- | -- |
| 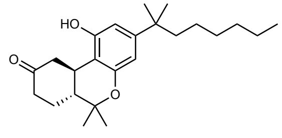 **Nabilone** | 2.3 [1.4 – 3.8]<br>8.6 [8.4 – 8.9]<br>81% [3.7] | -- | -- | -- | -- | -- |

N.D. = best fit values were not determined due to compound inactivity or poor data quality

-- Not tested

[a]One-way ANOVA statistical significance of individual comparisons to CP-55,940 after correction with Dunnett's test of multiple hypotheses. ns = not significant, * p<0.05, ** p<0.01, *** p<0.001, **** p<0.001

**Supplementary Table 3. Cryo-EM data collection, model refinement, and validation statistics.**

| **Data Collection** | **Global Refinement** |
|---|---|
| Voltage (kV) | 300 |
| Magnification | 96,000 |
| Total electron dose ($e^-/Å^2$) | 56.6 |
| Defocus range (μm) | -0.7 - -2.0 |
| Calibrated pixel size (Å) | 0.8521 |
| Micrographs collected | 8324 |
| **Data Processing** | |
| Extracted particles | 4,967,593 |
| Particles used for final reconstruction | 465,411 |
| Final map resolution (Å, 0.143 FSC) | 3.3 |
| Map resolution range (Å) | 2.6 - 4.2 |
| Map sharpening B factor ($Å^2$) | 175.3 |
| **Model Content** | |
| Initial models used (PDB code) | 6N4B (CB1/$G_i$/scFv16) |
| Total number of atoms | 8,528 |
| No. of protein residues | 1116 |
| No. of ligands | 1 |
| **Model Validation** | |
| CC map vs. model (%) | 69.05 |
| RMSD | |
| Bond lengths (Å) / Bond angles (°) | 0.003 / 0.663 |
| Ramachandran plot statistics | |
| Favored (%) | 94.12 |
| Allowed (%) | 5.88 |
| Outliers (%) | 0.0 |
| Rotamer outliers (%) | 0.0 |
| C-beta deviations | 0.0 |
| Clash score | 7.19 |

**Supplementary Table 4. Functional activities for select analogs versus a variety of transducers and hCB1 in the bioSens-All® platform.**

| Compound | | hCB1 $G_{i1}$ | hCB1 $G_{oB}$ | hCB1 $G_z$ |
|---|---|---|---|---|
| CP–55,940 | $EC_{50}$ [95% CI] (nM) | 0.46 [0.4 – 0.5] | 0.63 [0.6 – 0.7] | 0.28 [0.18 – 0.5] |
| | $pEC_{50}$ [95% CI] | 9.3 [9.3 – 9.4] | 9.2 [9.2 – 9.3] | 9.6 [9.4 – 9.8] |
| | $E_{max}$ [95% CI] | 100 [98 – 102] | 100 [99 – 101] | 102 [102 – 104] |
| '51486 | $EC_{50}$ [95% CI] (nM) | 849 [745– 947] | 711 [579– 873] | 1118 [363– 3447] |
| | $pEC_{50}$ [95% CI] | 6.1 [6.0 – 6.1] | 6.2 [6.1 – 6.2] | 6.0 [5.5 – 6.4] |
| | $E_{max}$ [95% CI] | 73 [70 – 75] | 74 [70 – 75] | 130 [996 – 163] |
| '60154 | $EC_{50}$ [95% CI] (nM) | 2.5 [1.9 – 3.4] | 18.4 [15.8– 21.5] | 17.5 [7.5 – 41] |
| | $pEC_{50}$ [95% CI] | 7.6 [7.5 – 7.7] | 7.7 [7.7 – 7.8] | 7.8 [7.4 – 8.1] |
| | $E_{max}$ [95% CI] | 92 [90 – 94] | 100 [98 – 100] | 121 [103 – 140] |
| '1081 | $EC_{50}$ [95% CI] (nM) | 150 [116 – 197] | 91.7 [55 – 153] | 225 [73 – 694] |
| | $pEC_{50}$ [95% CI] | 6.8 [6.7 – 6.9] | 7.0 [6.8 – 7.3] | 6.7 [6.2 – 7.1] |
| | $E_{max}$ [95% CI] | 49 [48 – 51] | 45 [43 –47] | 72 [55 –89] |
| '1082 | $EC_{50}$ [95% CI] (nM)<br>$pEC_{50}$ [95% CI]<br>$E_{max}$ [95% CI] | [a]N.D. | N.D. | N.D. |
| '1087 | $EC_{50}$ [95% CI] (nM)<br>$pEC_{50}$ [95% CI]<br>$E_{max}$ [95% CI] | N.D. | N.D. | N.D. |
| '1090 | $EC_{50}$ [95% CI] (nM) | 35.6 [28.5 – 45] | 37.3 [30 – 47] | 126 [45 – 354] |
| | $pEC_{50}$ [95% CI] | 7.5 [7.3 – 7.5] | 7.4 [7.3 – 7.5] | 6.9 [6.5 – 7.4] |
| | $E_{max}$ [95% CI] | 81 [80 –82] | 90 [88 –92] | 115 [92 – 137] |
| '4388 | $EC_{50}$ [95% CI] (nM)<br>$pEC_{50}$ [95% CI]<br>$E_{max}$ [95% CI] | N.D. | N.D. | N.D. |
| '6829 | $EC_{50}$ [95% CI] (nM) | 4056 [2417 – 9228] | 1011 [706 – 1449] | 1347 [720 – 2522] |
| | $pEC_{50}$ [95% CI] | 5.4 [5.0 – 5.6] | 6.0 [5.8 – 6.2] | 5.9 [6.0 – 6.2] |
| | $E_{max}$ [95% CI] | 68 [57 – 77] | 49 [45 – 53] | 70 [57 –83] |
| '4051 | $EC_{50}$ [95% CI] (nM) | 6523 [5770 – 7511] | 7988 [6709 – 9511] | 2431 [1069 – 5531] |
| | $pEC_{50}$ [95% CI] | 5.2 [5.1 – 5.2] | 5.1 [5.0 – 5.2] | 5.6 [5.3 – 6.0] |
| | $E_{max}$ [95% CI] | 108 [104 – 113] | 115 [106 – 121] | 114 [86 –141] |
| '12565 | $EC_{50}$ [95% CI] (nM) | 104 [79 – 138] | 44.8 [27 – 73] | 54.6 [22.1 – 135] |
| | $pEC_{50}$ [95% CI] | 7.0 [6.9 – 7.1] | 7.4 [7.1 – 7.6] | 7.3 [6.9 – 7.7] |
| | $E_{max}$ [95% CI] | 28 [27 – 28] | 25 [24 –26] | 51 [41 – 61] |
| '10010 | $EC_{50}$ [95% CI] (nM) | 814 [717 – 932] | 801 [582– 1102] | 1396 [506 – 3853] |
| | $pEC_{50}$ [95% CI] | 6.1 [6.0 – 6.1] | 6.1 [6.0 – 6.2] | 5.9 [5.4 – 6.3] |
| | $E_{max}$ [95% CI] | 74 [70 – 74] | 71 [67 – 75] | 144 [113 –175] |
| '6439 | $EC_{50}$ [95% CI] (nM) | 65.5 [56.8 – 755] | 60.9 [52.9 – 70] | 339 [96 – 1195] |
| | $pEC_{50}$ [95% CI] | 7.2 [7.1 – 7.2] | 7.2 [7.2 – 7.3] | 6.5 [5.9 – 7.0] |
| | $E_{max}$ [95% CI] | 95 [93 – 96] | 100 [99 – 101] | 162 [125 –199] |
| '6448 | $EC_{50}$ [95% CI] (nM) | 345 [321 – 372] | 310 [267 – 360] | 728 [381 – 1393] |
| | $pEC_{50}$ [95% CI] | 6.5 [6.4 – 6.5] | 6.5 [6.4 – 6.6] | 6.1 [5.9 – 6.4] |
| | $E_{max}$ [95% CI] | 87 [86 – 88] | 90 [87 – 91] | 151 [130 –172] |
| '3737a | $EC_{50}$ [95% CI] (nM) | 2804 [2436 – 3285] | 6729 [2678 – 16910] | 2470 [998 – 6108] |
| | $pEC_{50}$ [95% CI] | 5.6 [5.5 – 5.6] | 5.2 [4.8 – 5.6] | 5.6 [5.2 – 6.1] |
| | $E_{max}$ [95% CI] | 98 [94 – 102] | 118 [94 – 143] | 140 [9109 – 172] |
| '3737b | $EC_{50}$ [95% CI] (nM) | 3026 [2486 – 3822] | 19780 [4343 – 90080] | 30840 [1133 – 839000] |
| | $pEC_{50}$ [95% CI] | 5.5 [5.4 – 5.6] | 4.7 [4.1 – 5.4] | 4.5 [3.1– 6.0] |
| | $E_{max}$ [95% CI] | 89 [84 – 94] | 152 [94 – 208] | 231 [34 – 428] |
| '3737c | $EC_{50}$ [95% CI] (nM)<br>$pEC_{50}$ [95% CI]<br>$E_{max}$ [95% CI] | N.D. | N.D. | N.D. |
| '3737d | $EC_{50}$ [95% CI] (nM)<br>$pEC_{50}$ [95% CI]<br>$E_{max}$ [95% CI] | N.D. | N.D. | N.D. |

| | | | | |
|---|---|---|---|---|
| **‘7019** | **$EC_{50}$ [95% CI] (nM)**<br>**$pEC_{50}$ [95% CI]**<br>**$E_{max}$ [95% CI]** | N.D. | N.D. | [b]-- |
| **‘5424** | **$EC_{50}$ [95% CI] (nM)**<br>**$pEC_{50}$ [95% CI]**<br>**$E_{max}$ [95% CI]** | N.D. | N,D. | -- |
| **‘7800** | **$EC_{50}$ [95% CI] (nM)**<br>**$pEC_{50}$ [95% CI]**<br>**$E_{max}$ [95% CI]** | 6793 [983]<br>5.2 [6.0]<br>34 [24] | 597 [398 – 1017]<br>6.2 [5.9 – 6.4]<br>21 [20 – 23] | -- |
| **‘5463** | **$EC_{50}$ [95% CI] (nM)**<br>**$pEC_{50}$ [95% CI]**<br>**$E_{max}$ [95% CI]** | 4941 [1763 – 26860000]<br>5.3 [3.4 – 5.8]<br>31 [24 – 88] | 7472 [1125]<br>5.1 [6.0]<br>32 [22] | -- |
| **‘0450** | **$EC_{50}$ [95% CI] (nM)**<br>**$pEC_{50}$ [95% CI]**<br>**$E_{max}$ [95% CI]** | 56310 [2177]<br>4.3 [5.7]<br>49 [26] | 2509 [680 – 11020000]<br>5.6 [3.0 – 6.2]<br>36 [30 – 117] | -- |
| **‘2153** | **$EC_{50}$ [95% CI] (nM)**<br>**$pEC_{50}$ [95% CI]**<br>**$E_{max}$ [95% CI]** | 1011 [371 – 6576]<br>6.0 [5.2 – 6.4]<br>89 [78 – 118] | 2061 [620 – 25390]<br>5.7 4.6 – 6.2]<br>109 [91 – 162] | -- |

[a]N.D. = best fit values were not able to be determined due to compound inactivity or poor data quality
[b]-- Not tested

**Supplementary Table 5. Detailed functional activities for select analogs and controls versus a variety of transducers and hCB1 in the bioSens-All® platform.**

| Compound | | hCB1 $G_{i1}$ | hCB1 $G_{i2}$ | hCB1 $G_{oB}$ | hCB1 $G_z$ | hCB1 $G_{13}$ | hCB1 $G_{15}$ | hCB1 Barr2 + GRK2 |
|---|---|---|---|---|---|---|---|---|
| **CP–55,940** | **$EC_{50}$ [95% CI] (nM)** | 0.46 [0.4 – 0.5] | 0.55 [0.4 – 0.7] | 0.63 [0.6 – 0.7] | 0.28 [0.18 – 0.5] | 2.4 [1.65 – 3.4] | 0.26 [0.22 – 2.9] | 3.1 [1.97 – 4.8] |
| | **$pEC_{50}$ [95% CI]** | 9.34 [9.3 – 9.4] | 9.26 [9.1 – 9.4] | 9.20 [9.2 – 9.3] | 9.55 [9.4 – 9.8] | 8.62 [8.5 – 8.8] | 9.59 [9.5 – 9.7] | 8.51 [8.3 – 8.7] |
| | **$E_{max}$ [95% CI]** | 100 [98 – 102] | 100 [96 – 103] | 100 [98 – 101] | 101 [96 – 107] | 100 [95 – 105] | 100 [98 – 102] | 89 [82 – 98] |
| **'4042** | **$EC_{50}$ [95% CI] (nM)** | 0.48 [0.4 – 0.6] | 0.56 [0.4 – 0.9] | 0.64 [0.5 – 0.8] | 0.43 [0.3 – 0.6] | 2.1 [0.5 – 9.6] | 0.37 [0.28 – 0.49] | 3.6 [2.1 – 6.8] |
| | **$pEC_{50}$ [95% CI]** | 9.32 [9.2 – 9.4] | 9.25 [9.0 – 9.5] | 9.20 [9.1 – 9.3] | 9.37 [9.3 – 9.4] | 8.69 [8.0 – 9.3] | 9.43 [9.3 – 9.6] | 8.44 [8.2 – 8.7] |
| | **$E_{max}$ [95% CI]** | 102 [100 – 104] | 101 [95 – 106] | 103 [99 – 106] | 97 [93 – 102] | 64 [54 – 84] | 105 [102 – 108] | 72 [71 – 74] |
| **'1350** | **$EC_{50}$ [95% CI] (nM)** | 0.23 [0.18 – 0.3] | 0.28 [0.19 – 0.4] | 0.29 [0.24 – 0.34] | 0.35 [0.22 – 0.53] | 0.83 [0.23 – 2.7] | 0.22 [0.18 – 0.27] | 2.2 [1.1 – 4.4] |
| | **$pEC_{50}$ [95% CI]** | 9.63 [9.5 – 9.7] | 9.54 [9.4 – 9.7] | 9.54 [9.5 – 9.6] | 9.46 [9.3 – 9.7] | 9.08 [8.6 – 9.6] | 9.66 [9.6 – 9.8] | 8.66 [8.4 – 9.0] |
| | **$E_{max}$ [95% CI]** | 92 [90 – 95] | 95 [90 – 99] | 98 [94 – 98] | 94 [88 – 100] | 56 [48 – 67] | 99 [96 – 101] | 63 [60 – 72] |
| **'8690** | **$EC_{50}$ [95% CI] (nM)** | 19.1 [14.4 – 25.8] | | 18 [14 – 23] | 33 [15 – 117] | | | |
| | **$pEC_{50}$ [95% CI]** | 7.72 [7.6 – 7.8] | [b]-- | 7.8 [7.7 – 7.9] | 7.5 [6.9 – 7.8] | -- | -- | [a]N.D. |
| | **$E_{max}$ [95% CI]** | 82 [77 – 85] | | 83 [80 – 87] | 98 [86 – 121] | | | |
| **2-AG** | **$EC_{50}$ [95% CI] (nM)** | 224 [147 – 353] | 62 [42 – 918] | 93 [65 – 135] | 493 [15] | 4470 [1618 – 398200] | 32 [25 – 40] | 1025 [774 –1446] |
| | **$pEC_{50}$ [95% CI]** | 6.65 [6.5 – 6.8] | 7.2 [7.0 – 7.4] | 7.0 [6.9 – 7.2] | 6.3 [7.8] | 5.4 [5.1 – 5.8] | 7.5 [7.4 – 7.6] | 6.0 [5.8 – 6.1] |
| | **$E_{max}$ [95% CI]** | 122 [113 – 133] | 112 [106 – 119] | 114 [108 – 121] | 205 [112] | 183 [133 – 607] | 112 [108 – 116] | 200 [184 – 220] |

[a]N.D. = best fit values were not able to be determined due to compound inactivity or poor data quality
[b]-- Not tested

**Supplementary Table 6. Relative efficacy calculations for '4042 and '1350 versus CP-55,940.**

| Target | Sensor | Compound | Mean log ($E_{max}/EC_{50}$) | SEM log ($E_{max}/EC_{50}$) | Mean Δlog ($E_{max}/EC_{50}$) | SEM Δlog ($E_{max}/EC_{50}$) | t-test to CP-55,940[a] | RE[b] |
|---|---|---|---|---|---|---|---|---|
| hCB1 | $G_{i1}$ | **CP-55,940** | 9.34 | 0.07 | 0.00 | 0.09 | | 1.00 |
| | | **'4042** | 9.33 | 0.14 | -0.01 | 0.15 | ns | 0.97 |
| | | **'1350** | 9.59 | 0.06 | 0.30 | 0.09 | **** | 1.98 |
| | $G_{i2}$ | **CP-55,940** | 9.24 | 0.13 | 0.00 | 0.18 | | 1.00 |
| | | **'4042** | 9.25 | 0.03 | 0.01 | 0.13 | ns | 1.01 |
| | | **'1350** | 9.52 | 0.03 | 0.28 | 0.13 | ** | 1.89 |
| | $G_{oB}$ | **CP-55,940** | 9.19 | 0.07 | 0.00 | 0.10 | | 1.00 |
| | | **'4042** | 9.21 | 0.09 | 0.02 | 0.11 | ns | 1.04 |
| | | **'1350** | 9.09 | 0.44 | 0.35 | 0.44 | **** | 2.24 |
| | $G_z$ | **CP-55,940** | 9.53 | 0.23 | 0.00 | 0.33 | | 1.00 |
| | | **'4042** | 9.32 | 0.19 | -0.21 | 0.30 | ns | 0.62 |
| | | **'1350** | 9.42 | 0.22 | -0.11 | 0.32 | ns | 0.77 |
| | $G_{13}$ | **CP-55,940** | 8.63 | 0.23 | 0.00 | 0.32 | | 1.00 |
| | | **'4042** | 8.59 | 0.25 | -0.04 | 0.33 | ns | 0.91 |
| | | **'1350** | 8.85 | 0.05 | 0.22 | 0.23 | * | 1.64 |
| | $G_{15}$ | **CP-55,940** | 9.59 | 0.02 | 0.00 | 0.02 | | 1.00 |
| | | **'4042** | 9.46 | 0.12 | -0.13 | 0.12 | ns | 0.74 |
| | | **'1350** | 9.65 | 0.06 | 0.06 | 0.06 | ns | 1.14 |
| | $ß_{arr2}$+ GRK2 | **CP-55,940** | 8.28 | 0.25 | 0.00 | 0.35 | | 1.00 |
| | | **'4042** | 8.19 | 0.03 | -0.09 | 0.03 | * | 0.81 |
| | | **'1350** | 8.34 | 0.08 | 0.06 | 0.08 | ns | 1.16 |
| hCB2 | $G_{i1}$ | **CP-55,940** | 8.86 | 0.12 | 0.00 | 0.17 | | 1.00 |
| | | **'4042** | 8.49 | 0.03 | -0.20 | 0.13 | ns | 0.63 |
| | | **'1350** | 8.32 | 0.10 | -0.54 | 0.16 | * | 0.29 |
| | $G_{i2}$ | **CP-55,940** | 8.97 | 0.00 | 0.00 | 0.00 | | 1.00 |
| | | **'4042** | 8.74 | 0.00 | -0.23 | 0.00 | [c]-- | 0.59 |
| | | **'1350** | 8.57 | 0.00 | -0.40 | 0.00 | -- | 0.40 |
| | $G_{oB}$ | **CP-55,940** | 8.76 | 0.09 | 0.00 | 0.13 | | 1.00 |
| | | **'4042** | 8.43 | 0.19 | -0.33 | 0.21 | n.s. | 0.47 |
| | | **'1350** | 8.36 | 0.01 | -0.40 | 0.09 | *** | 0.40 |
| | $G_z$ | **CP-55,940** | 8.90 | 0.39 | 0.00 | 0.55 | | 1.00 |
| | | **'4042** | 8.48 | 0.51 | -0.42 | 0.64 | n.s. | 0.38 |
| | | **'1350** | 8.24 | 0.23 | -0.65 | 0.45 | n.s. | 0.22 |
| | $ß_{arr2}$+ GRK2 | **CP-55,940** | 7.83 | 0.13 | 0.00 | 0.18 | | 1.00 |
| | | **'4042** | 7.74 | 0.00 | -0.09 | 0.13 | *** | 0.82 |
| | | **'1350** | 7.79 | 0.09 | -0.04 | 0.16 | n.s. | 0.92 |

[a]Statistical significance of all comparisons of compound activities (Mean Δlog ($E_{max}/EC_{50}$)) to CP-55,940 control by unpaired t-test. ns = not significant, * $p<0.05$, ** $p<0.01$, *** $p<0.001$, **** $p<0.0001$

[b]RE, relative efficacy = $10^{\Delta\log (Emax/EC50)}$

[c]--, not determined

### Supplementary Table 7. Binding affinities and functional activities for select active analogs at CB2.

| Compound | rCB2 binding<br>$EC_{50}$ [95% CI] (nM)<br>$pEC_{50}$ [95% CI]<br>$E_{max}$ [SEM] | hCB2<br>Cerep cAMP<br>$EC_{50}$ [95% CI] (nM)<br>$pEC_{50}$ [95% CI]<br>$E_{max}$ [95% CI] | hCB2<br>BRET2 + GoA<br>$EC_{50}$ [95% CI] (nM)<br>$pEC_{50}$ [95% CI]<br>$E_{max}$ [95% CI] | hCB2<br>Tango<br>ß-arrestin<br>recruitment<br>$EC_{50}$ [95% CI] (nM)<br>$pEC_{50}$ [95% CI]<br>$E_{max}$ [95% CI] |
|---|---|---|---|---|
| 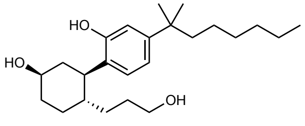<br>**CP–55,940** | -- | 0.082 | 13.1 [9.4 – 18.5]<br>7.9 [7.7 – 8.0]<br>99% [94 – 105] | 22.9 [21.3 – 24.8]<br>7.6 [7.60 – 7.67]<br>100% [98 – 102] |
| 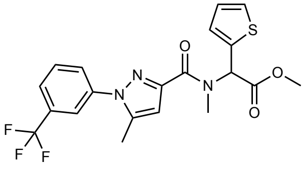<br>**'4042** | 2.18 [1.7 – 2.8]<br>8.7 [8.6 – 8.8]<br>103% [8.8] | 0.011 [0.002 – 0.02]<br>10.95 [10.7 – 11.6]<br>72% [77 – 84] | 7.9 [1.6 – 40.6]<br>8.1 [7.4 – 8.8]<br>29% [22 – 36] | 33.9 [21.7 – 61.5]<br>7.5 [7.2 – 7.7]<br>28% [26 – 32] |
| 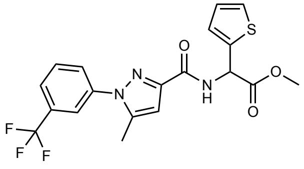<br>**'60154** | [b]-- | -- | 594 [125 – 2813]<br>6.2 [5.6 – 6.9]<br>31% [20 – 42] | 3341 [2707 – 4371]<br>5.5 [5.4 – 5.6]<br>21% [19 – 23] |
| 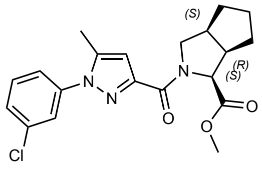<br>**10010** | -- | -- | [a]N.D. | 556 [506 – 611]<br>6.3 [6.2 – 6.3]<br>18% [17 – 19] |

| | | | | |
|---|---|---|---|---|
| 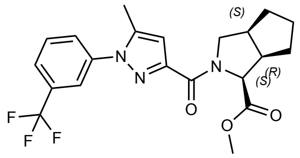 **'6439** | -- | -- | 97.1 [12.6 – 749]<br>7.0 [6.13 – 67.9]<br>81% [34 – 128] | 338 [282– 415]<br>6.5 [6.4 – 6.6]<br>14% [13 – 15] |
| 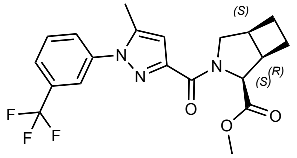 **'6448** | -- | -- | -- | -- |
| 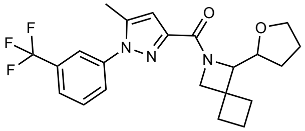 **'3737** | -- | 4.3 [.0004 – 22.6]<br>8.4 [7.7 – 12.5]<br>45% [42 – 80%] | N.D. | 1500 [1274 – 1805]<br>5.8 [5.7 – 5.9]<br>9% [8 – 9] |

[a]N.D. = best fit values were not determined due to compound inactivity or poor data quality
[b]-- Not tested

**Supplementary Table 8. Functional activities for select analogs and controls versus a variety of transducers and hCB2 in the bioSensAll platform.**

| Compound | | hCB2 $G_{i1}$ | hCB2 $G_{i2}$ | hCB2 $G_{oB}$ | hCB2 $G_z$ | hCB2 Barr2 + GRK2 |
|---|---|---|---|---|---|---|
| **CP–55,940** | **$EC_{50}$ [95% CI] (nM)** | 1.4 [1.1 – 1.7] | 1.1 [0.9 – 1.3] | 1.7 [1.5 – 2.1] | 1.05 [0.6 – 1.9] | 14.3 [11.6 – 17.9] |
| | **$pEC_{50}$ [95% CI]** | 8.87 [8.7 – 8.9] | 8.97 [8.9 – 9.1] | 8.76 [8.7 – 8.8] | 8.98 [8.7 – 9.2] | 7.84 [7.8 – 7.9] |
| | **$E_{max}$ [95% CI]** | 100 [97 – 103] | 100 [97 – 104] | 100 [98 – 103] | 98 [89 – 109] | 100 [95 – 104] |
| **‘4042** | **$EC_{50}$ [95% CI] (nM)** | 2.7 [2.0 – 3.4] | 1.3 [0.9 – 1.7] | 2.5 [1.9 – 3.3] | 1.36 [0.5 – 3.1] | 5.5 [3.5 – 8.7] |
| | **$pEC_{50}$ [95% CI]** | 8.58 [8.5 – 8.7] | 8.90 [8.8 – 9.0] | 8.60 [8.5 – 8.7] | 8.87 [8.5 – 9.3] | 8.26 [8.1 – 8.5] |
| | **$E_{max}$ [95% CI]** | 82 [79 – 85] | 70 [67 – 73] | 61 [59 – 63] | 56 [47 – 68] | 33 [33 – 33] |
| **‘1350** | **$EC_{50}$ [95% CI] (nM)** | 3.55 [2.98 – 4.2] | 1.6 [1.4 – 1.9] | 2.6 [2.15 – 3.1] | 2.6 [1.6 – 3.9] | 4.2 [2.8 – 6.3] |
| | **$pEC_{50}$ [95% CI]** | 8.45 [8.4 – 8.5] | 8.79 [8.7 – 8.9] | 8.58 [8.5 – 8.7] | 8.58 [8.4 – 8.8] | 8.38 [8.2 – 8.6] |
| | **$E_{max}$ [95% CI]** | 74 [72 – 77] | 61 [59 –63] | 59 [57 – 61] | 52 [46 – 60] | 30 [28 –33] |
| **2-AG** | **$EC_{50}$ [95% CI] (nM)** | 217 [186 – 256] | 96 [70 – 129] | 394 [349 – 447] | 2123 [931 – 87070] | 1854 [1110–4061] |
| | **$pEC_{50}$ [95% CI]** | 6.66 [6.6 – 6.7] | 7.0 [6.9 – 7.2] | 6.4 [6.3 – 6.5] | 5.7 [4.1 – 6.0] | 5.7 [5.4 – 6.0] |
| | **$E_{max}$ [95% CI]** | 105 [102 – 109] | 78 [74 – 84] | 101 [98 – 104] | 96 [76 – 233] | 118 [104 – 140] |
